## Supplementary Materials & Methods for "A Hierarchical 3D-motion Learning Framework for Animal Spontaneous Behavior Mapping"

1. Shenzhen Key Lab of Neuropsychiatric Modulation and Collaborative Innovation Center for Brain Science, Guangdong Provincial Key Laboratory of Brain Connectome and Behavior, CAS Center for Excellence in Brain Science and Intelligence Technology, Brain Cognition and Brain Disease Institute (BCBDI), Shenzhen Institutes of Advanced Technology, Chinese Academy of Sciences, Shenzhen-Hong Kong Institute of Brain Science-Shenzhen Fundamental Research Institutions, Shenzhen 518055, China.
2. University of the Chinese Academy of Sciences, Beijing 100049, China.
3. Pennsylvania State University, State College, Pennsylvania, 16802, USA.
4. Co-first authors
5. Lead Contact

#### Abstract:

Animal behavior usually has a hierarchical structure and dynamics. Therefore, to understand how the neural system coordinates with behaviors, neuroscientists need a quantitative description of the hierarchical dynamics of different behaviors. However, the recent end-to-end machine-learning-based methods for behavior analysis mostly focus on recognizing behavioral identities on a static timescale or based on limited observations. These approaches usually lose rich dynamic information on cross-scale behaviors. Inspired by the natural structure of animal behaviors, we addressed this challenge by proposing a novel parallel and multi-layered framework to learn the hierarchical dynamics and generate an objective metric to map the behavior into the feature space. In addition, we characterized the animal 3D kinematics with our low-cost and efficient multi-view 3D animal motion-capture system. Finally, we demonstrated that this framework could monitor spontaneous behavior and automatically identify the behavioral phenotypes of the transgenic animal disease model. The extensive experiment results suggest that our framework has a wide range of applications, including animal disease model phenotyping and the relationships modeling between the neural circuits and behavior.

**Key Words:** Behavioral structure-inspired; 3D motion capture; Behavioral dynamics; Computational ethology; Behavior phenotyping.

### Supplementary Figures

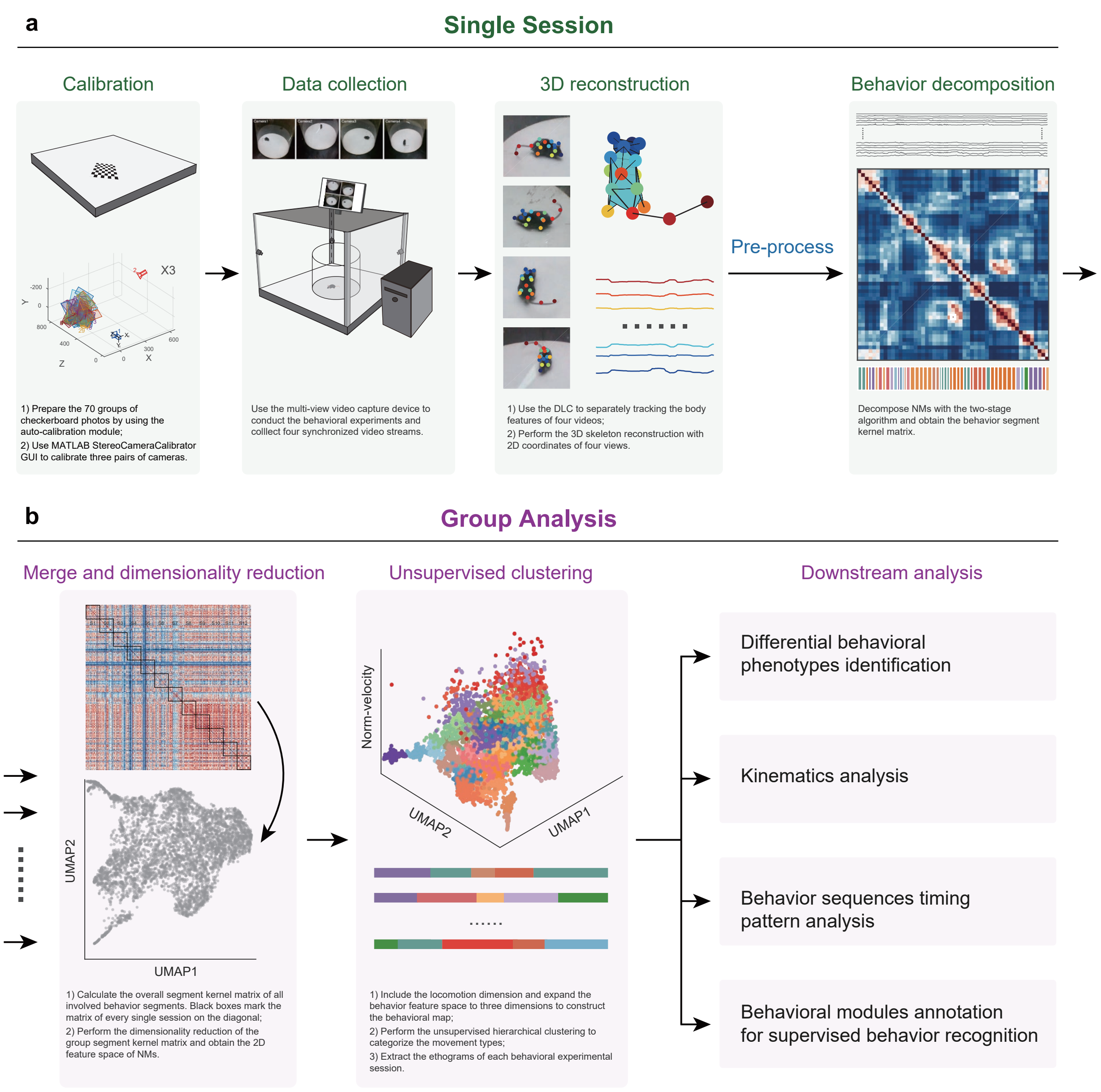


**Supplementary Fig. 1 | The workflow of the hierarchical 3D-motion learning framework. a** Four main steps for a single experimental session: 1) Calibration (related to Supplementary Methods "The calibration of 3D motion capture system"). Using the auto-calibration module to quickly prepare 70 groups of checkerboard images from various angles and positions for calibration and using the MATLAB StereoCameraCalibrator GUI to calculate the calibration parameters of the three pairs of cameras. This step is necessary only when the calibration parameters are unknown, or the cameras have been moved. 2) Data collection (related to Supplementary Methods Animals, behavioral experiments and behavioral data collection). Setting up the behavioral apparatus and preparing the animal and then using the multi-view video capture device to collect the synchronous behavioral videos. 3) 3D reconstruction (related to Supplementary Methods 3D pose reconstruction). Using the DLC pre-trained model to predict the animal's 16 body-part 2D coordinates from the four separate videos, then performing the 3D skeleton reconstruction with the 2D coordinates from four views to obtain the animal's postural time-series. 4) Behavior decomposition (related to Supplementary Methods Behavior decomposition). Performing the two-stage behavior decomposition on the pre-processed postural time-series. This step discovers the behavioral modules based on the optimal movement segmentation. Finally, these behavioral segments are aligned using the DTAK metric to construct the segment kernel matrix representing their similarity. **b** Group analysis based on specific biological questions. 1) Merge and dimensionality reduction (related to Supplementary Methods: Group segment kernel matrix and low dimensional embedding). According to experimental grouping, single session segment kernel matrices are merged into a group segment kernel matrix. To visualize the informative structure of the behavioral modules involved, we used dimensionality reduction to transform the group segment kernel matrix into a 2D space. 2) Unsupervised clustering (related to Supplementary Methods: Unsupervised clustering). Constructing the behavioral map by combining the NM space with the locomotion dimension, then using the unsupervised clustering algorithm to categorize the movement sequence into distinct types. After clustering, ethograms can be constructed by associating the behavioral labels with their original segments. 3) Downstream analysis. After obtaining each session's ethogram, the downstream quantitative analysis can be conducted according to experimental grouping, recording stage, and other conditions to answer biological questions from behavioral aspects.

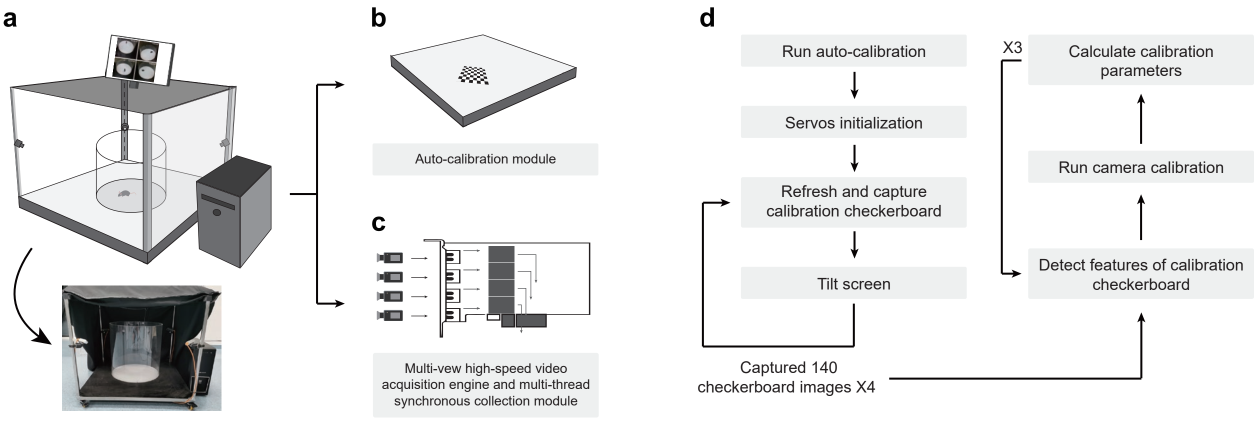


**Supplementary Fig. 2 | Illustration of the multi-view video capture device and the workflow of the auto-calibration module. a** Schematic of the multi-view video capture device. The support framework is a 90 × 90 × 75 $cm^3$ movable stainless steel shelf, on which cameras, behavioral apparatus, calibration modules, and background lighting are mounted. A shielding curtain can be added per experimental requirements. **b** The auto-calibration module is designed for efficient camera calibration in 3D and is composed of an LCD screen for displaying the checkerboard and a control unit used to tilt the screen. To collect images of the checkerboard pattern at different orientations relative to the cameras, the calibration program controls the screen to rotate and translate the checkerboard pattern at different tilt angles. With this auto-calibration module, the checkerboard images can be captured in one minute. **c** The multi-view video acquisition module. Four video streams, one per camera, are input to the PCI-E USB-3.0 data acquisition card (expanded bandwidth). The acquisition program then uses multi-thread acquisition to ensure frame synchronization. **d** The two-part workflow of the auto-calibration program. The first part, shown on the left, automatically collects a variety of checkerboard patterns for each camera (70). Right, the calibration process, which is based on the MATLAB StereoCameraCalibrator GUI.

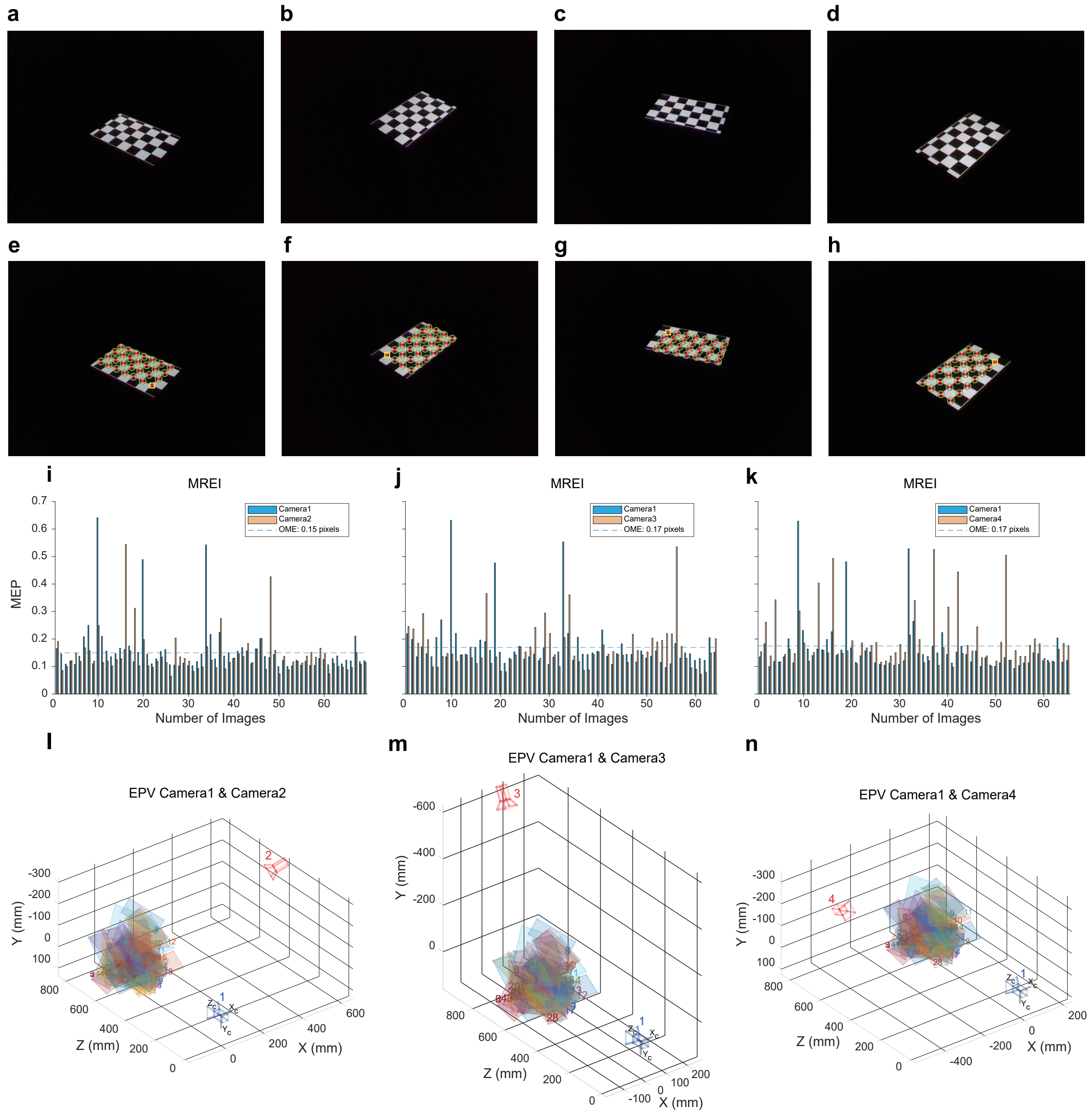


**Supplementary Fig. 3 | Calibration of cameras results. a-d** one of 70 checkerboard images of each camera in four different placements. All of the checkerboards are fully captured in four different cameras placements with dark light. We use a 10-inch tablet to display checkerboard to calibrate four cameras, which could freely adjust the brightness and get appropriate brightness easily to keep the checkerboard clear enough. **e-h** the result of grid detection with MATLAB's StereoCameraCalibrator GUI corresponding to **a-c** and **d** images. The corner recognition of the checkerboard is located on the corner of the black squares. **i-k** mean reprojection of error per image (MREI). MREI gives the quantization level of calibration error between cameras and checkerboards, where the mean error in pixels (MEP) represents the pixel errors in each camera-checkerboard pair and overall mean error (OME) gives a whole estimation of pixel errors. **l-n** extrinsic parameters visualization (EPV) of the cameras. EPV shows the relative positions of two cameras and checkerboards in 3D space. If EPV is different from real relative positions, we should calibrate two cameras again until they are coincident.

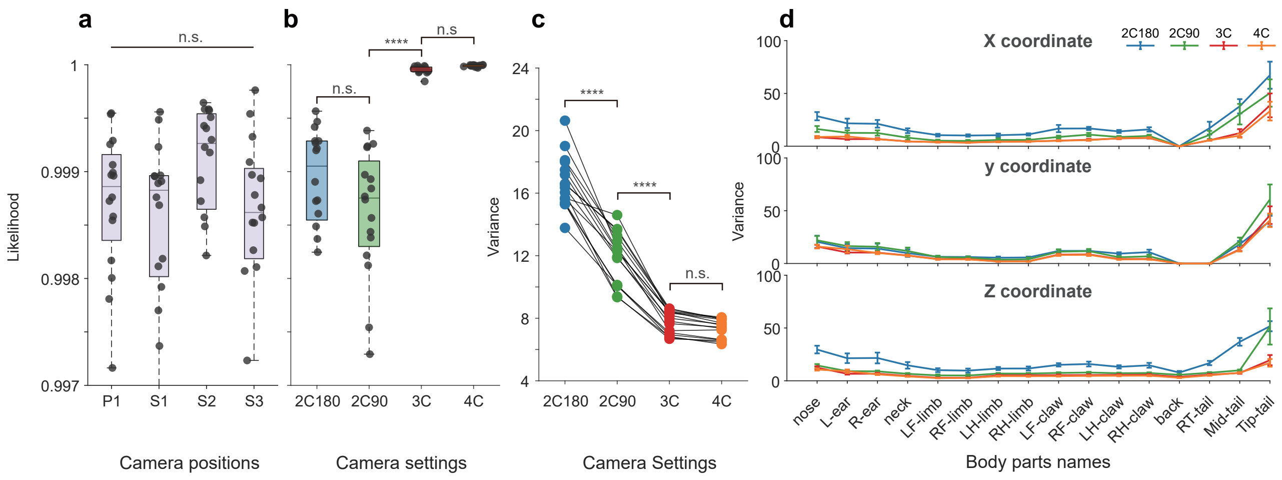


**Supplementary Fig. 4 | Evaluation of 3D Reconstruction Quality with Different Camera Settings.** **a** The likelihoods of the DLC pose estimations of four camera positions. P1, primary camera 1, S1, secondary camera 1, S2, secondary camera 2, S3, secondary camera 3. Each point on the boxplot represents the mean likelihood of each test recording, which is calculated by firstly averaging the likelihoods of all the body parts per frame then averaging them across all frames. The likelihoods show no significant differences among these cameras (Kruskal-Wallis test, $p = 0.1339$, $n = 16$). **b** The likelihoods of the 3D reconstructions of different camera groupings. 2C180, two cameras are placed in opposite directions. 2C90, two cameras are positioned in orthogonal directions. 3C, three cameras. 4C, four cameras. In the camera groupings 2C180 and 2C90, each point on the boxplot is calculated by firstly specifically averaging the likelihoods of two paired body parts for calibration, then averaging all 16 paired averaged likelihoods per frame and finally averaging them across all frames. In the camera groupings of 3C and 4C, each point on the boxplot is calculated by firstly specifically averaging the first two maximum likelihoods of paired body parts for calibration from all the three or four points, then averaging all 16 paired averaged maximum likelihoods per frame, and finally averaging them across all frames. **c** The variances of the behavioral trajectories captured by different camera groupings. Each point on the plot is calculated by firstly computing the variances of each body part's trajectory in the X, Y, Z axis, then averaging them across X, Y, Z axis, and finally averaging them across 16 body parts. (Kruskal-Wallis test, ****$p < 0.0001$, $n = 16$). **d** The variances of each body part in X, Y, Z coordinates of varying camera groupings. The variances of each body part are calculated by firstly computing the variances of each body part's trajectory in X, Y, Z axis then averaging them across X, Y, Z axis.

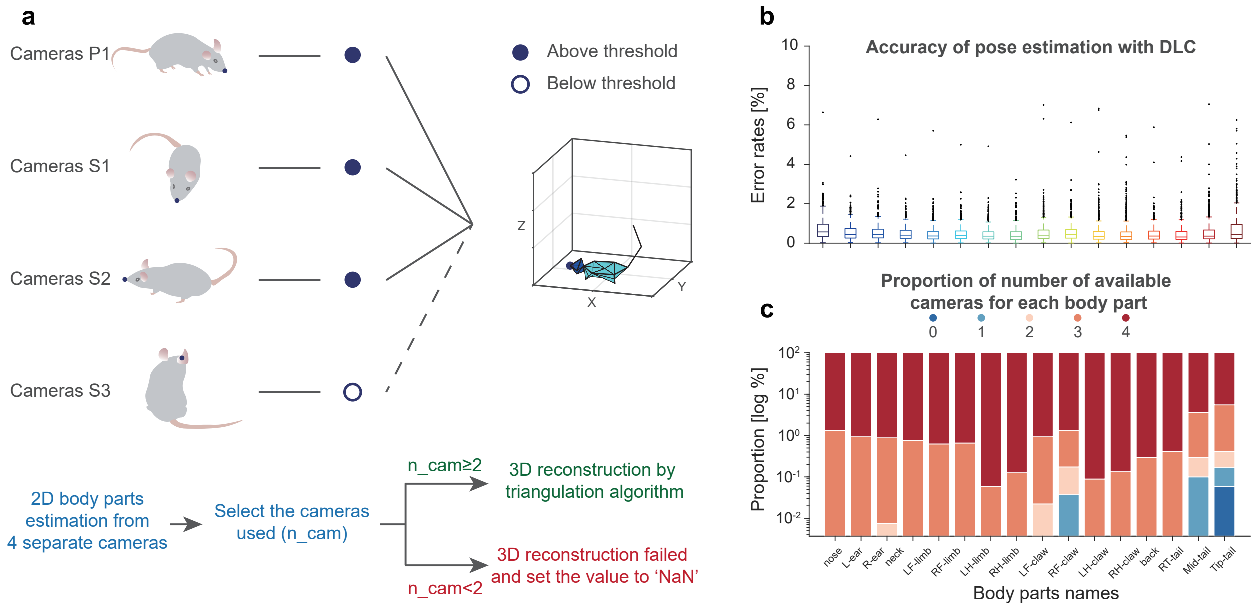


**Supplementary Fig. 5 | 3D reconstruction process and reliability evaluation of the occluded body parts. a** The workflow of 3D reconstruction of a single body part: 1) estimate the two-dimensional coordinates of the animal's body part from four cameras; 2) select the cameras to be used for reconstruction by thresholding the likelihood of the estimated body part; 3) determine whether the number of cameras available meets the reconstruction requirements (2 or more); and 4) if 2 or more, reconstruct the 3D coordinate of the body part. Otherwise, the 3D reconstruction fails due to the occlusion. P1, primary camera. S1, first secondary camera. S2, second secondary camera. S3, third secondary camera. **b** The errors in 2D body-part estimations versus ground truth. The error rates are shown for each body part separately in the boxplot and averaged 0.534 ± 0.005%. **c** The proportional number of available cameras by body part. The average proportions are: no cameras, 0.004%; 1 camera, 0.015%; 2 cameras, 0.038%; 3 cameras, 1.048%; 4 cameras, 98.895%.

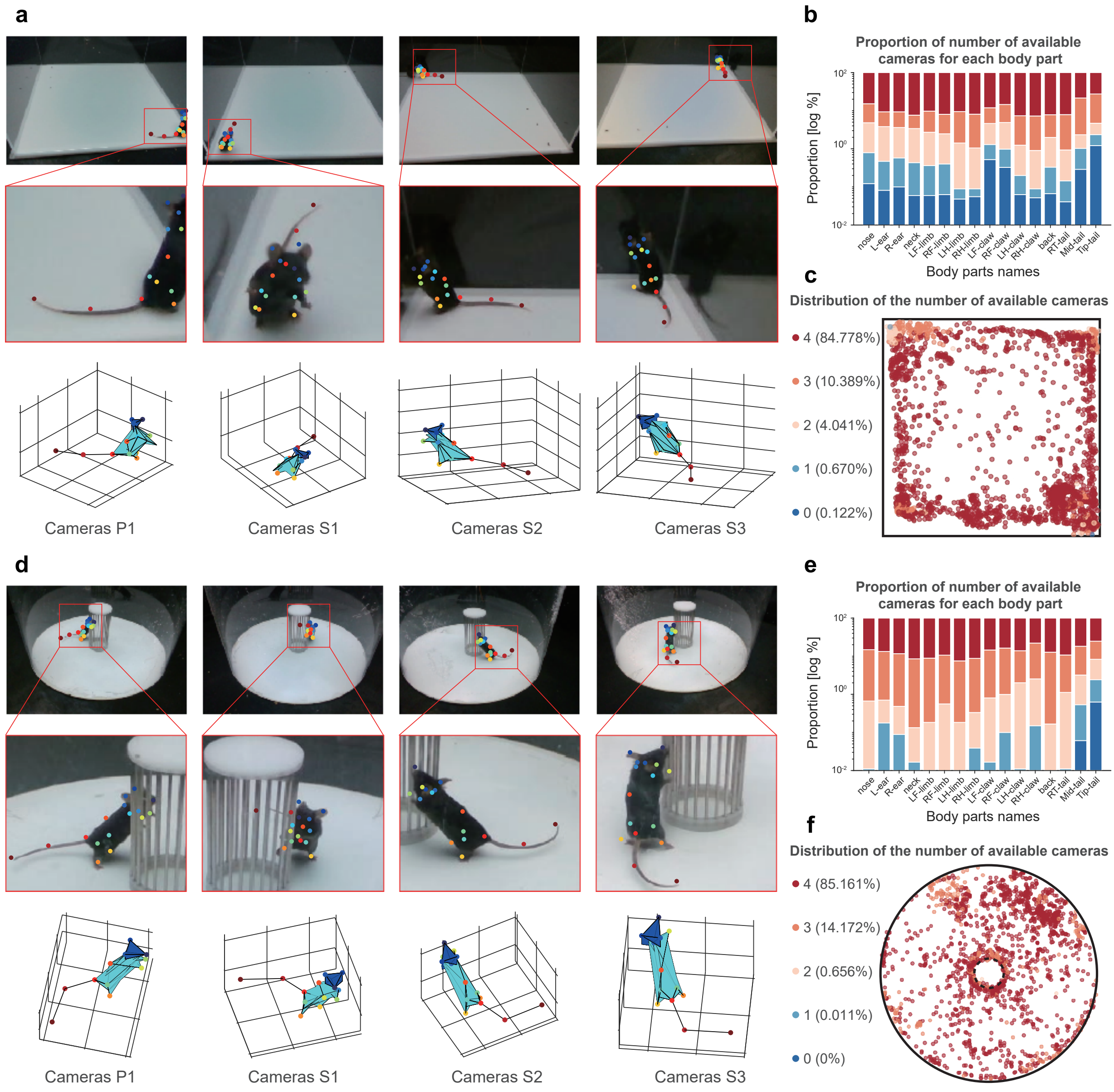


**Supplementary Fig. 6 | Evaluation of the 3D reconstruction in cases of view-point–specific disappearances of body parts. a**, **d** 2D pose tracking and 3D skeleton reconstruction of representative view-point disappearance frames from two different test apparatuses. **a** First test: square open field test. The behavior chamber is out of the field of view, and blind areas may occur when the animal enters the four corners; **d** Second test: circular open-field with a sociability cage. The mouse can easily be occluded by the cage, thus blind areas may exist in one or more perspectives. Top: selected frames with one or more views in which body parts disappear. Middle: magnification to show the disappearance details. Bottom: successfully reconstructed 3D skeletons shown in approximately the same views as the corresponding recordings. **b**, **e** The proportional number of cameras available for 3D reconstruction for each body part. In the first test, an average of 99.398 ± 0.149% of all frames showing the body part can meet the reconstruction requirements; In the second test, the average reconstruction rate of all body parts is 99.776 ± 0.150%. **c**, **f** The distribution of the number of available cameras for 3D reconstruction.

The color-coded dots indicate how many cameras are used for reconstruction at the indicated location. The positions of all the points are the x and y coordinates of the nose. For visualization purposes, the data are down-sampled to 10%. The proportional number of cameras used to reconstruct the nose in 3D is indicated in the key on the left.

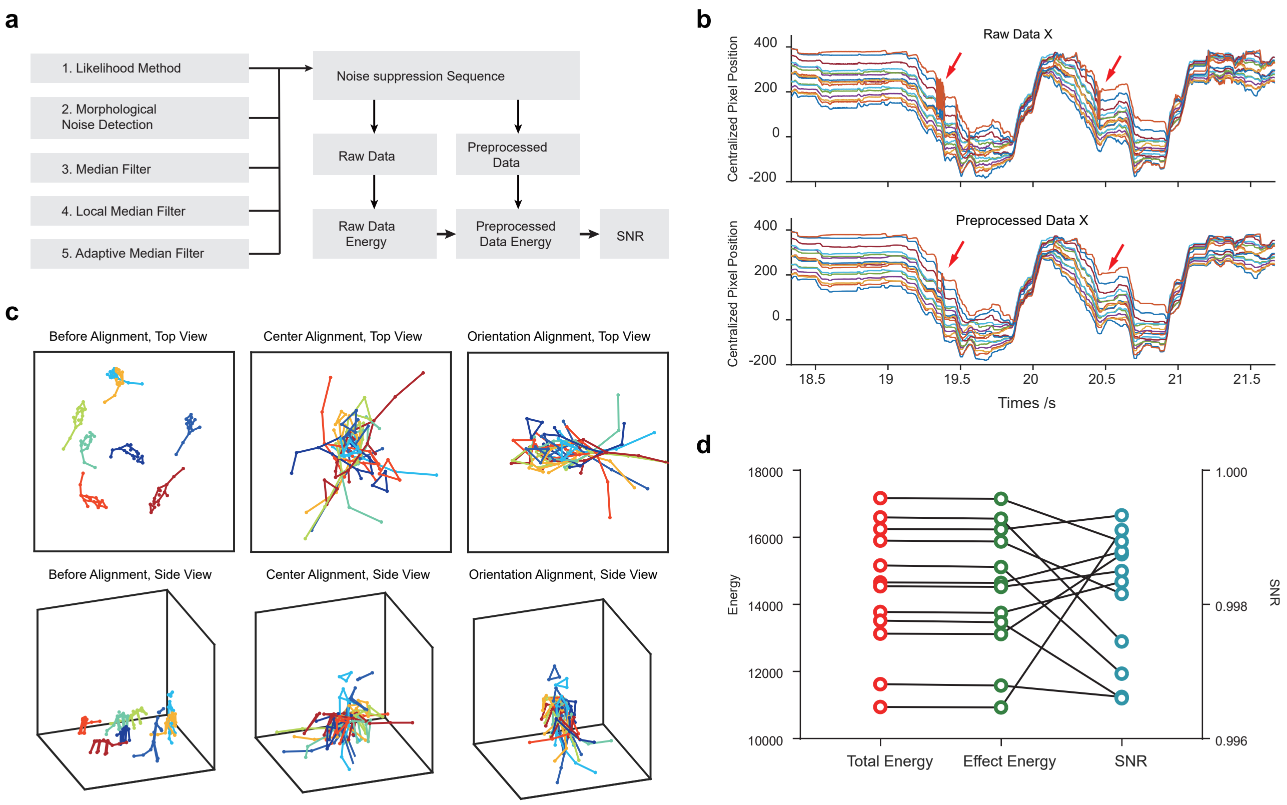


**Supplementary Fig. 7 | The procedure of data preprocessing. a** The procedure of data quality control contains two main parts, a noise suppression sequence part ordered by five filtering modules and the SNR detection part. **b** The selected multi-dimensional time series of mouse skeleton are preprocessed by a data quality control procedure. The red arrows indicate that the apparent noises are suppressed, and the whole series doesn't miss too much rapidly changing details. **c** Eight skeleton frames are randomly selected for a demo of mice alignment. The results of center orientation alignment are demonstrated on the top view and side view. **d** The multi-dimensional time-series data quality of 12 involved test mice are described, whose SNRs are more than 0.996, representing that there are averaged only four noise points in a thousand data points needed to be eliminated.

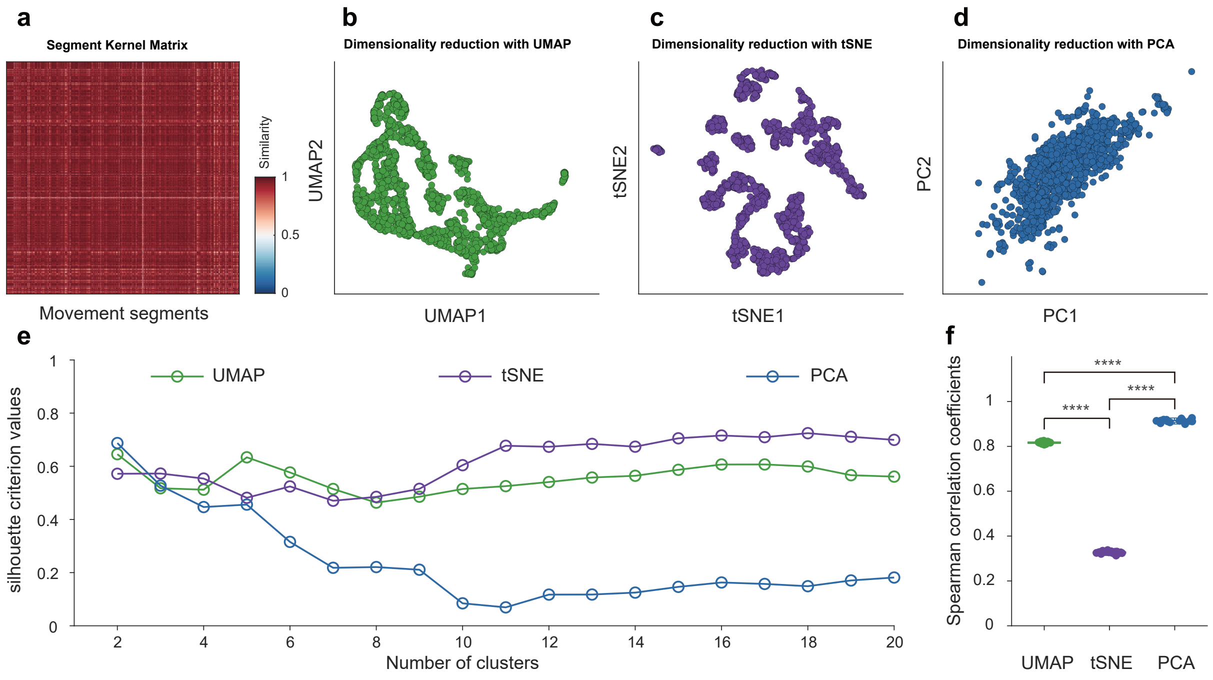


**Supplementary Fig. 8 | Comparison of three algorithms of dimensionality reduction for the representation of the NM feature space structure. a** Segment kernel matrix of a representative single-session behavioral experiment recording. The matrix pixels represent the normalized similarity value of 937 pairs of decomposed movement segments. **b-d** Dimensionality reduction with the three most-used algorithms: UMAP, tSNE (t-distributed stochastic neighbor embedding), and PCA (principal component analysis). For visualization purposes, the segment kernel matrix is reduced to two dimensions. **e** Quantification of local structure preservation by evaluation of the silhouette criterion values of the dimensionality reduction result of each algorithm. The silhouette criterion values are calculated by enumerating the clusters from two to twenty. The average silhouette criterion values are: UMAP, 0.557 ± 0.011; tSNE, 0.619 ± 0.021; PCA, 0.240 ± 0.039. **f** Quantification of global structure preservation by evaluation of the Spearman correlation coefficients between the original segment kernel matrix and the dimensionality-reducted result of each algorithm. For each algorithm, we first randomly subsampled 70% of the kernel matrix 20 times. Each time, the Spearman correlation coefficients are calculated between the selected segment kernel sub-matrix and the paired-wise distances of the dimensionality-reduced data. The average coefficients are: UMAP, 0.817 ± 0.001; tSNE, 0.326 ± 0.001; PCA, 0.913 ± 0.002. ****, P < 0001 by two-way ANOVA with a Holm–Sidak post-hoc test.

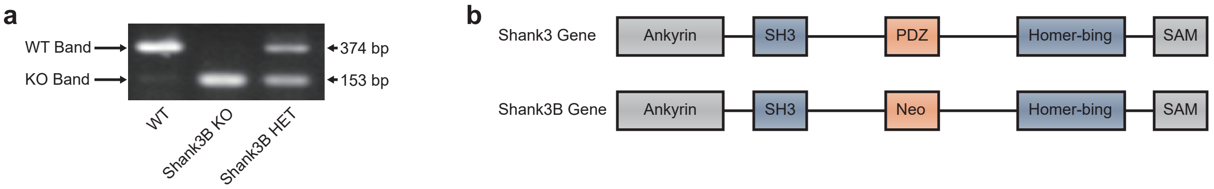


**Supplementary Fig. 9 | Genotyping for *Shank3* mice. a** PCR genotyping for *Shank3B*$^{+/+}$(WT), *Shank3B*$^{-/-}$ (Shank3B KO), and *Shank3B*$^{+/-}$ (Shank3B HET) mice. **b** working strategy for *Shank3B*-mutant mice.

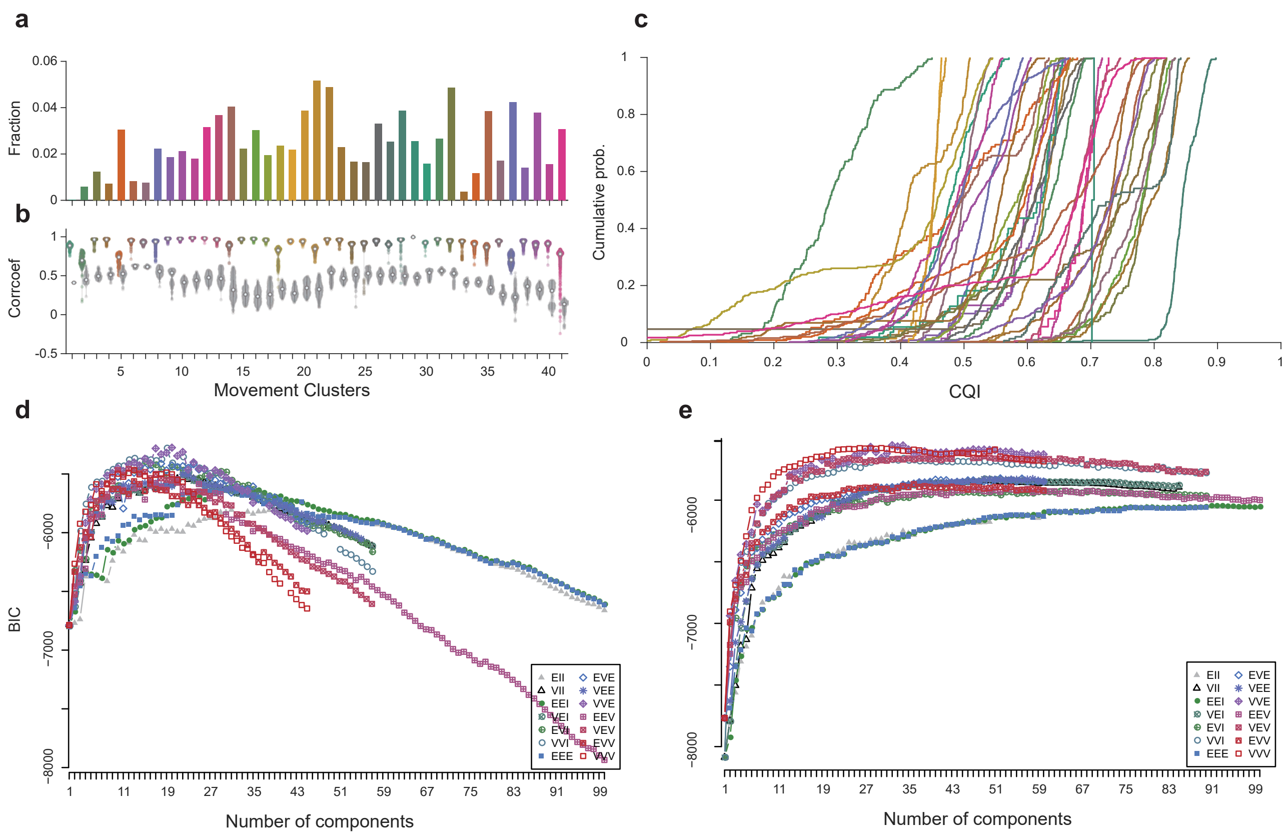


**Supplementary Fig. 10 | Clustering quality evaluation and the number of clusters determination.** **a** the fractions of movement bouts number (total number: 16607), the color-coded bars indicate the clustered movement types (totally 41 types). **b** the intra-CC (color-coded), and inter-CC (grey dots) of each movement group. The dots on each violin plot represents their intra-CC or inter-CC, and dots number in a pair of violin plot in each group are the same. **c** the cumulative distribution function of CQI of the movement clusters. The clusters represented by the curves on the right side have better clustering qualities, and their corresponding movements are more stereotyped. **d** the BIC of single-session experiment shows (related to Fig. 4) the most appropriate number of clusters in movement clustering could be chosen in the range of 10 to 20. **e** the BIC of all the mice's movements (related to Fig. 6) in movement space shows when the number of clusters is beyond about 20, the BIC of all the movements is tending to be steady, and the maximum BIC is in the range of 35 to 45.

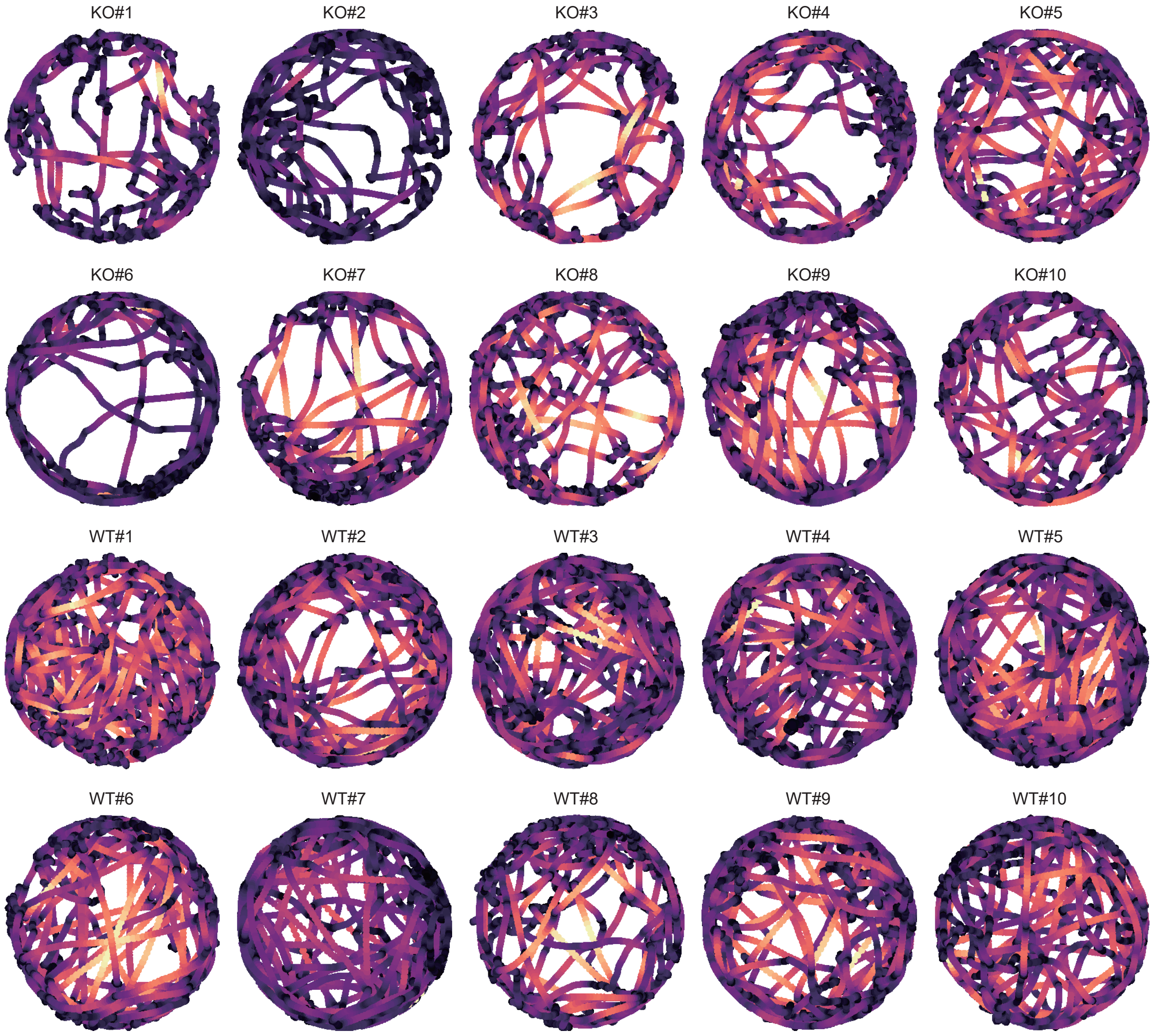


**Supplementary Fig. 11 | The velocity-trajectory heatmaps of all the 10 *Shank3B*$^{-/-}$ (KO) mice and 10 *Shank3B*$^{+/+}$ (WT) mice.** Compare the velocity-trajectory heatmaps between the two groups, KO mice have lower activity levels than WT mice.

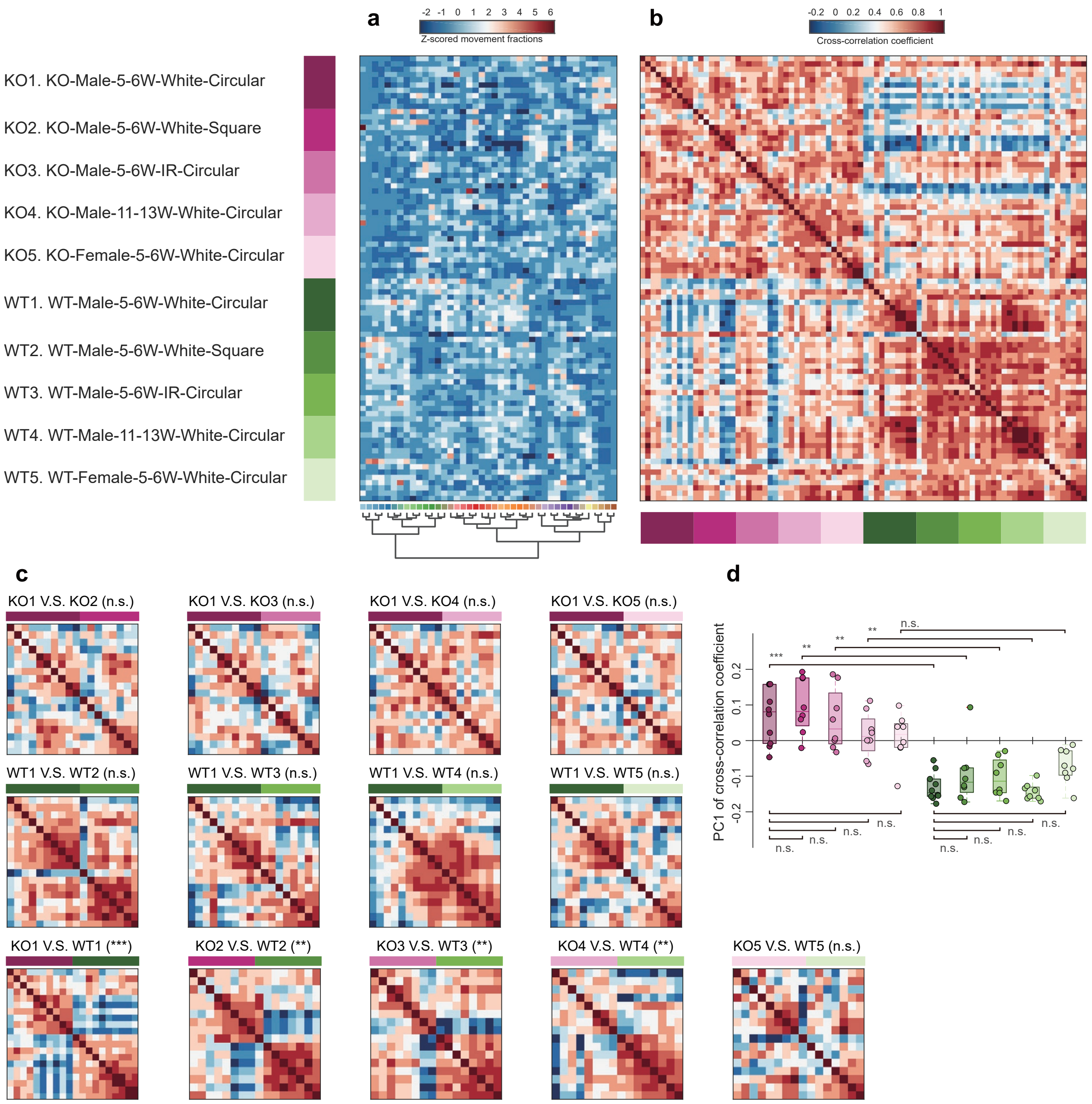


**Supplementary Fig. 12 | Group comparison of Shank3B KO mice under different conditions. a** The movement fraction matrix of mice in ten different groups. The color-bars shown in the left indicate the group conditions for the corresponding rows of the matrix (see Supplementary Methods for further details). Each row in the fraction matrix represents the tested mouse, and each column corresponds to the behavioral module types arranged with the dendrogram in the bottom. For visualization and comparison purpose, the values of movement fraction matrix are normalized with z-score by rows. In each group, the row orders are determined by placing the sample with largest variance of the movement fraction, and then the other samples are ranked according to the decreasing correlation with the first row. **b** The cross-correlation coefficients matrix (CCCM) of the movement fractions among all ten groups samples. **c** The group comparisons of behavioral correlations between the selected conditions, which are shown with twelve submatrices of **b**. **d** The behavioral statistics between ten groups. The comparison metric is determined by calculating the principal component (PC) of the CCCM, then using the first PC (PC1) to evaluate the overall behavioral differences across ten groups (Kruskal-Wallis test, **, $p<0.01$, ***, $p<0.001$).

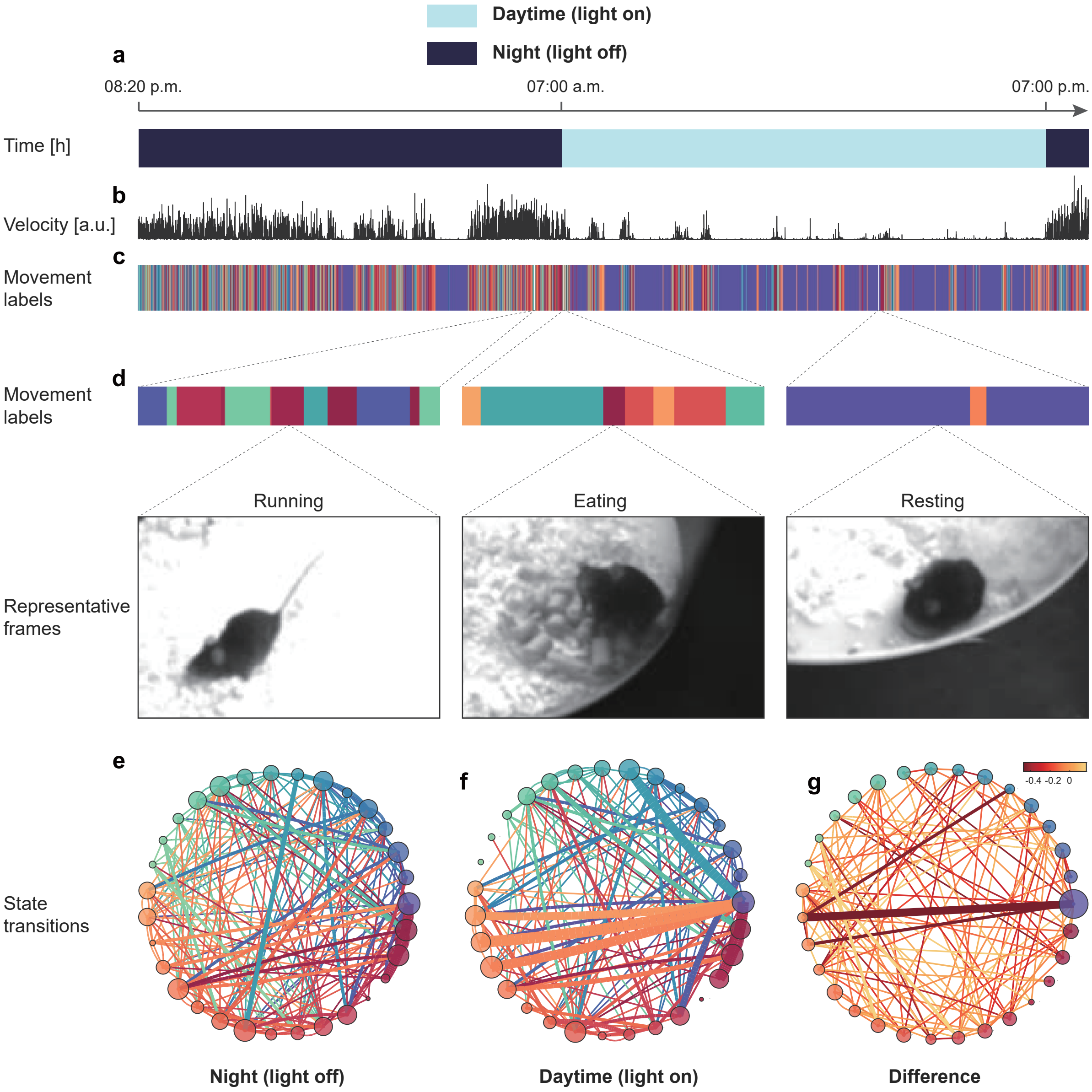


**Supplementary Fig. 13 | Continuous long-term monitoring and analysis of mouse behavior. a** The timeline of the behavioral recording period over 24 hours. **b** The normalized velocity of the mouse across 24 hours aligned to the timeline. **c** The decomposed behavioral modules shown with color-coded labels. **d** Three magnified representative behavioral modules and selected, single corresponding frames. Left, running on the litter; Middle, eating; Right, prolonged immobility resembling resting. **e, f** State transitions of the movement modules in night and day phases. **g** Differences in the state transitions between night and day. The color of the dots in **e, f,** and **g** correspond to the behavioral modules shown in **c**. The size of the dots represents the rank of the module probabilities over 24 hours. The color of the connections in **e** and **f** represents the direction from the previous state to the current state, and its color is the same as that of the previous state. The width of the connections in **e** and **f** represents the normalized two-state transition probability. The color and width of the connections in **g** represent the normalized difference between **e** and **f**.

### Supplementary Methods

#### Apparatus

The multi-view video capture device shown in Supplementary Fig. 1a and Supplementary Fig. 2a, b. Mice freely walk in a 50 cm diameter and 50 cm height circular open field [1] with transparent acrylic wall and white plastic ground. The circular open field is placed on the center of a 90 × 90 × 75 $cm^3$ movable stainless steel support framework. A black thick, dull polish rubber mat is paved between the circular open field and steel shelf to avoid light reflection. The four supporting pillars of the shelf are mounted four Intel RealSense D435 cameras [2], and cameras are placed orthogonally. Images are simultaneously recorded at 30 frames per second by PCI-E USB-3.0 data acquisition card using the pyrealsense2 Python camera interface package. On the top of the shelf, a 56-inch television was facing down and placed horizontally to provide uniform and stable white background light. The cameras and TV are connected to a high-performance computer (i7-9700K, 16G RAM) equipped with a 1-terabyte SSD and a 12-terabyte HDD as a platform for the software and hardware of images acquisition.

#### The calibration of 3D motion capture system

We use MATLAB's StereoCameraCalibrator GUI to finish the cameras' calibration of 3D motion capture system. The GUI uses the basic principles of Zhang's calibration method [3], and we need to use some checkerboard pictures to obtain parameters of 2D to 3D.

Firstly, we use the code of Python and OpenCV to control four cameras capturing checkerboard pictures synchronously. We have two different checkerboard display schemes in all of our behavioral experiments. The old version is using a 10-inch tablet to display checkerboard. The experimenter needs to control the synchronously images capturing of four cameras by a keyboard shortcut then move the checkerboard to different positions until each camera captures 70 images of checkerboard (Supplementary Fig. 3a-d). To improve the efficiency of checkerboard capturing step, we update the 3D motion capture system. The new version changes the bottom surface of the system by a square 32-inch displayer. We merge the code of checkerboard display, checkerboard moving and four cameras checkerboard capturing together to achieve the automatic checkerboard images capturing by 3D motion capture system. In the process of automatic camera calibration debugging, we found that only displaying the checkerboard in a horizontal plane would confuse the optimizing of camera parameters by StereoCameraCalibrator GUI. Hence, we developed a mechanical device like a table concentrator to tilt the square displayer to make sure the success rate of camera parameters optimizing.

After the step of checkerboard images capturing, the parameters of cameras could be calculated by them. We assume that X, Y, and Z are 3D coordinate points of stereo cameras so that the equation can be written as

$$P\begin{pmatrix} u \\ v \\ 1 \end{pmatrix} = \begin{pmatrix} \alpha & \gamma & u_0 \\ 0 & \beta & v_0 \\ 0 & 0 & 1 \end{pmatrix} \begin{pmatrix} \mathbf{R} & \mathbf{t} \end{pmatrix} \begin{pmatrix} X \\ Y \\ Z \\ 1 \end{pmatrix} \quad (1)$$

where $P$ is scaling constants, $u$ and $v$ are pixel coordinates of the checkerboard image, $\alpha$ is the focal length divided by the length of the picture, $\beta$ is the focal length divided by the width of the picture, $\gamma$ is radial distortion parameter, $u_0$ and $v_0$ are midpoint in pixel coordinates of the image, $\mathbf{R}$ is a rotation matrix, $\mathbf{t}$ is translation vector. Zhang's calibration method can obtain $\alpha$, $\beta$, $\gamma$, $\mathbf{R}$, $\mathbf{t}$, $P$. As the previous statement, each camera takes 70 calibration photos, and we can get 280 calibration photos using four cameras (Supplementary Fig. 3). Meanwhile, we measure and record the grid's size of the checkerboard. We set one of the cameras as the primary camera, and the remaining three cameras serve as sub-cameras. The cameras are divided into three groups, and each group contains a secondary camera and a primary camera (see Supplementary Fig. 4). In each group, we use StereoCameraCalibrator GUI to calibrate two cameras and can get three calibrated files.

**Animals, behavioral experiments and behavioral data collection**

Young adult (5-6 weeks) male, normal adult (11-13 weeks) male, young adult (5-6 weeks) female *Shank3B*$^{-/-}$ and *Shank3B*$^{+/+}$ mice with C57BL/6J genetic background were used in the behavioral experiment (Supplementary Fig. 9, and Supplementary Table 2). *Shank3B*$^{-/-}$ mice were obtained from the Jackson Laboratory (Jax No. 017688). *Shank3B*$^{-/-}$ and *Shank3B*$^{+/+}$ mouse lines have been described previously [4] . The mice were housed at 4–6 mice per cage under a 12-h light-dark cycle at 22–25°C, and were allowed to acquire water and food ad libitum. All husbandry and experimental procedures in this study were approved by Animal Care and Use Committees at the Shenzhen Institute of Advanced Technology (SIAT), Chinese Academy of Sciences (CAS). All behavioral experiments were performed and analyzed blinded to genotypes.

We designed two behavioral experiments. In the first behavioral experiment, we collected the *Shank3B*$^{-/-}$ and *Shank3B*$^{+/+}$ mice behavioral data under different conditions (experimental apparatus, lighting conditions, ages, and genders). There are total ten different groups (see Supplementary Fig. 12, and Supplementary Table 2), and respectively are: 1) KO1: genotype *Shank3B*$^{-/-}$, age 5-6 weeks, male, circular open field, white light; 2) KO2: genotype *Shank3B*$^{-/-}$, age 5-6 weeks, male, square open field, white light; 3) KO3: genotype *Shank3B*$^{-/-}$, age 5-6 weeks, male, circular open field, infrared light; 4) KO4: genotype *Shank3B*$^{-/-}$, age 11-13 weeks, male, circular open field, white light; 5) KO5: genotype *Shank3B*$^{-/-}$, age 5-6 weeks, female, circular open field, white light; 6) WT1: genotype *Shank3B*$^{+/+}$, aged 5-6 weeks, male, circular open field, white light; 7) WT2: genotype *Shank3B*$^{+/+}$, age 5-6 weeks, male, square open field, white light; 8) WT3: genotype *Shank3B*$^{+/+}$, age 5-6 weeks, male, circular open field, infrared light; 9) WT4: genotype *Shank3B*$^{+/+}$, aged 11-13 weeks, male, circular open field, white light; 10) WT5: genotype *Shank3B*$^{+/+}$, aged 5-6 weeks, female, circular open field, white light. Each mouse in each group is only used once. In the 10 groups above, mice were allowed to explore the open field for 10 minutes freely, then 15-minute records were analyzed. We write codes of Python and OpenCV to obtain and record videos of mice. The frame rate is set to 30 fps and the frame size is set to 640×480.

In the second behavioral experiment, we capture the mouse behavior for 24 hours (related to Supplementary Fig. 13). We still use circular open field but covered by wood chip as padding and offered water and regular food (the chow). The male mouse used in this experiment has C57BL/6J genetic background and is 13 weeks old. To change the light conditions and keep the circadian rhythms of mouse, we use infrared light as the background light and set the cameras to infrared model. This experiment was start at 20:20 p.m. We first closed the light until 7:00 a.m. next day, then we open the white light until 19:00 p.m. At last, we closed the light until 20:20 p.m. and finished the behavioral capturing across 24 hours. All the detailed information of mice and experimental conditions are in Supplementary Table 2.

**Mouse pose estimation**

The DeepLabCut (DLC) toolbox[7] is used to track the animal's two-dimensional (2D) body features from four recorded separate videos. We manually labeled about 3000 images as a training set to improve the environment adaptation of DLC pose estimation model. The maximum number of iterations is 1,200,000, and the final cross-entropy loss is 0.017. The training step spends 20 hours on NVIDIA GTX 2080Ti GPU. We use the trained network model to track new video and obtain 16 feature points at each frame of a video, including the nose, neck, and so on. Each video produces one feature file, and we can obtain four feature files per trail of experiments.

**3D pose reconstruction**

We use pose3d toolbox in MATLAB to finish 3D reconstruction of the animal skeleton. Pose3d toolbox uses the triangulation algorithm to obtain 3D data, and triangulation equation is

$$\mathbf{x}_i = \mathbf{P}_i\mathbf{X} \tag{2}$$

where $\mathbf{x}_i = [m_i, n_i, 1]^T$ , $\mathbf{X}=[I, J, K, 1]^T$ , and $\mathbf{P}_i = \mathbf{A}_i[\mathbf{R}_i, \mathbf{t}_i]$ . $\mathbf{x}_i$ is the coordinate of the matching point in the $i$ camera image, $\mathbf{X}$ is a vector of 3D point, $I$ , $J$ , $K$ are the 3D point of $\mathbf{X}$ , $\mathbf{P}_i$ is the projection matrix, $\mathbf{A}_i$ is the internal reference of the $i^{th}$ camera, $\mathbf{R}_i$ is the rotation matrix of the $i^{th}$ camera, $\mathbf{t}_i$ is the $i^{th}$ translation vector of the $i^{th}$ camera. Using the least square method to get $\mathbf{X}$ , and the solution of the least square method is obtained by singular value decomposition (SVD) [8]. Specifically, three calibrated files and four feature files are inputted to the system of pose3d, and the system produces one 3D feature files as output. After this, we need to rotate and translate 3D coordinates to the horizon for visualization and later processing. Using the Delaunay triangulation [9] to fill the skeleton, which makes it look much more like a live mouse.

**Evaluation of 3D Reconstruction Quality with Different Camera Settings**

To test the limit of reducing the number of cameras, we demonstrate detail analysis of precision quantification with different camera settings (Supplementary Fig. 4). For each camera, 2D pose estimation's likelihoods have no significant difference, which means the position of the camera has no significant impact on the estimation result. In the 3D reconstruction procedure, it is enough to only apply two calibrated cameras for the acquisition of 3D body points. So, the best two points with the highest likelihoods from all four cameras are selected to be reconstructed in the 3D space. We choose four different cameras groupings (2C180, 2C90, 3C, 4C) to verify reconstruction accuracy (Fig. 2b). The likelihoods of 2C180 and 2C90 have no significant differences, which means that in the case of two cameras, the position differences do not affect the mouse's basic 3D reconstruction. The likelihoods of 3C and 4C also have no significant differences, meaning that three cameras could basically reach the precision requirement of four cameras in this case. Both 3C and 4C's likelihoods are significantly higher than the 2C180 and 2C90 group (Kruskal-Wallis test, ****$p<0.0001$, $n=16$). This result corresponds with multiple previous studies, showing that the amount of camera has a positive correlation with the accuracy of pose-tracking [10, 11]. From this point of view, the minimum number of cameras is two. But for less occlusion, the results and previous studies suggest to increase the number of cameras focusing on different animals.

Besides that, we calculate the 3D behavior trajectories' variances of different camera settings. Variance is a representation of data information. The information of data has two parts, objective information, and noise information. The statistics analysis shows that 2C180 is significantly higher than 2C90, and 2C90 is markedly higher than 3C. But 3C and 4C have no significant difference. According to the proportional relations of them, we could calculate that 2C180 has more than 30% noise information compared with 2C90, and 2C90 has more than 50% noise information compared with 3C and 4C. Thus, the best and economic number of cameras is three in this experiment situation. When the variance is converged with more cameras, we could find the least number of cameras to reduce the cost. Therefore, according to the analysis of trajectories' variances of different camera settings, we could test the limit of reducing the number of cameras.

At last, we test the influence of accuracy of each body part in difference camera settings. We calculated the variances of each body part in each X, Y and Z coordinate of 3D poses. Different camera settings have the same trend in the variances of different body points. Significantly, the variance of tails in 3D poses is higher than other body parts. These results suggest that for different body points, the noise level and tracking ability of same camera settings are different. Tracking specific body points need balance the number, quality, and grouping of cameras. Such as the tail of mice, if the movement of tail is the crucial research object, it is important to add more cameras to improve the tracking accuracy.

**The proportion of the number of available cameras for each body part**

In the 3D reconstruction step, it has the probability that not all of the 2D body points in multi-view cameras could be used for 3D reconstruction because of tracking failure caused by the occlusion (Supplementary Fig. 5, 6). For this reason, we evaluate the accuracy of 3D reconstruction from the view of the number of available cameras for each body part. A threshold of the likelihood of pose estimation is used to judge if the tracking of specific body points is failure. If the likelihood is larger than the threshold, the camera is used. The threshold is set to 0.9, and we use a video of 27000 frames for the calculation of the proportion. The number of cameras to use is showed by scaled stack bars.

**Data pre-processing**

Data preprocessing have two procedures.

The first procedure is the data quality control (Supplementary Fig. 7a). In order to ensure the precision of follow-up algorithms, the data quality needs to be controlled. Data quality control has two steps.

The first step is data noise suppression. We build a suppression algorithm sequence of data noise, including likelihood method, morphological noise detection, median filter [12], local median filter, and adaptive median filter, which would be introduced in detail later. These algorithms could be ordered as needed.

The likelihood method is a noise detection step. The likelihood of raw data in DLC is counted, and a threshold is set to distinguish noise level likelihood and non-noise level likelihood. We set the threshold as 95% when the likelihood is smaller than 95%, the noise data points are detected.

Morphological noise detection finds the leap of body points in the physical morphological level. The most precise pose estimation frame is selected automatically by the maximum likelihood frame as a template. The maximum body points distance of template multiplied a coefficient is set as a threshold to detect morphological saltation of animal skeletons by comparing current maximum body points distance with the threshold. Then the distances between two points, which are higher than the threshold, would be marked as noise.

The median filter is a widely used method for impulse noise. It replaces the impulse noise points by median values in a time window, which could effectively eliminate the impulse noise while keeping the high-frequency details of data. Most of the noise data points in pose estimation could be regarded as impulse noise so that it is suitable for the median filter to suppress them.

The local median filter is used as an interpolation algorithm in our framework. We put the median filter window across the noise data points detected by the likelihood method and morphological filter, then replacing the noise data points by median data in the filter window. Because of the filter only focusing on noise location, it would not influence qualified data.

The adaptive median filter is designed to substitute the median filter. The median filter has a fixed window width so that it treats all data at the same time scale. The adaptive median filter adopts variable window width; thus, the impulse noise in different time scales could be disposed of more properly.

We provide a default parameter of noise suppression algorithms sequence for preprocessing our 3D animal pose estimation data (likelihood method, morphological noise detection, and median filter with 1 s time window).

The second step is the data quality assessment (Supplementary Fig. 7d). We calculate the multi-dimensional time series total energy of raw data, preprocessed data, and noise defined as the absolute value of the subtraction of raw data and preprocessed data. Then we use these energies to assigned a signal to noise ratio (SNR). The division of preprocessed data calculates SNR's total energy to raw data total energy.

The second procedure for data preprocessing is mice alignment (Supplementary Fig. 7c). The alignment has four steps, and all the alignments have proceeded in a 3D Cartesian coordinate system $\mathbf{\Theta} = \{\mathbf{X}, \mathbf{Y}, \mathbf{Z}\}$ .

The first step is center alignment. We regard the back point as a center point and make each value of body part subtract center point in the plane $\{\mathbf{X}, \mathbf{Y}\}$ of respectively. Each frame of mice skeletons would have the same center coordinate value (0, 0) after center alignment.

The second step is orientation alignment. We regard the vector from the back point to the tail root point as the orientation vector of mice. Thus, aligning the orientation of mice is the same as aligning the orientation of the orientation vector. We compute rotation matrixes from the orientation vector, and the positive direction of the $\mathbf{X}$ axis of each frame then rotates the mice skeleton to the same orientation by rotation matrixes in the plane $\{\mathbf{X}, \mathbf{Y}\}$. The data points in $\mathbf{Z}$ are not be changed, which retain the height information of mice skeletons.

The third step is size normalization. We find that the size of mice would cause a bias in poses and movements decomposition because of the weights of body size is always larger than body movements in different mice. Considering that the length of the orientation vector could represent the size of mice skeletons, we calculate the distribution of orientation vector lengths of each mice and choose the median value as the length pattern. Then we set a standard length (practical value from our tests of the range of orientation vector acquired by our apparatus, which is set to 25) and calculate a list of size normalization factors by the division of the median value of orientation vector length with the standard length. All the frames of skeletons would multiply by the factors in different mice to correct the mice size to standard size.

After the procedure of mice alignment, there are some dimensions of time series that could be reduced. The center points series in $\{\mathbf{X}, \mathbf{Y}\}$ , and the endpoint of orientation vectors in $Y$ axis are all zeros so that they would not influence the decomposition precision but the time consumption, which could be reduced undoubtedly. Besides that, we focus on the body movements of mice. Previous reports suggested that the movement of the tail is relatively independent from the rest of the body[13]. Considering that rodents also emit other specific signals corresponding to other body parts, with the exception of the torso, through the tail's movements, the following motion quantification did not involve the motion features of two parts of the tail. As our framework measures the inherent dynamic similarity of the time-series, the choice of features depends on the behavior of interest. Therefore, we reduce nine dimensions from 48 dimensions' mice time series and leave 39 dimensions for poses and movements decomposition.

**Behavior decomposition**

Poses and non-locomotor movements (NM) don't have locomotion features, which could be decomposed from the time series after alignment. Let $\mathbf{X} = \{\mathbf{x}_1, \mathbf{x}_2, ..., \mathbf{x}_{39}\}^T$ be the 39-dimension time series after alignment, where each time series $\mathbf{x}_i = (x_i^1, x_i^2, ..., x_i^N)^T$ contains $1 \times N$ ordered real values. Firstly the $\mathbf{X}$ is reduced spatial dimensions from 39 dimensions to 2 dimensions by uniform manifold approximation and projection (UMAP) [14], which could keep the global and local structures of data at the same time. Let $\mathbf{Z} = \{\mathbf{z}_1, \mathbf{z}_2\}^T$ be the dimension reducing time series and $f_{UMAP}$ be the mapping of $\mathbf{X}$ to $\mathbf{Z}$, then we get

$$\mathbf{Z} = f_{UMAP}(\mathbf{X}) \tag{3}$$

where $f_{UMAP}(\cdot)$ have the parameters n_neighbors to 80 and min_dist to 0.3.

Secondly, we use the temporal reduction to decompose poses from $\mathbf{Z}$ . The temporal reduction is using density clustering in the time dimension to cluster $\mathbf{Z}$ into $k_{poses}$ classes. The time dimension is represented by the distribution whose different centers of density could be regarded as different poses in $\mathbf{A}$ . We use default parameters of the density clustering method in UMAP to get the results of poses decomposition. Let $\mathbf{U} = \{\mathbf{u}_1, \mathbf{u}_2\}^T$ be the poses decomposition time series, $\mathbf{l} = \{l_1, l_2, .., l_N\}^T$ be the classes label and $f_{DC}$ be the density clustering mapping of $\mathbf{Z}$ to $\mathbf{U}$ , then we get

$$\mathbf{U} = f_{DC}(\mathbf{Z}) \tag{4}$$

where $\mathbf{U} \in \mathrm{IR}^{2\times M}$. $M$ is the number of poses, which is calculated by the temporal reduction length $r_L$ and continuous poses frames $\mathbf{Z}_{[p,p+L_p]}$, $p \in [1,M]$, where $L$ is the number of continuous poses frames. Each $\mathbf{Z}_{[p,p+L_p]}$ is decomposed into $\mathbf{Z}_{[p,p+l_p^j]}$ by $r_L$ and $l$ could be calculated as

$$l_p^j = \begin{cases} r_L, j < \mathrm{ceil}(\frac{L_p}{r_L}) \\ \mathrm{rem}(L_p, r_L), j = \mathrm{ceil}(\frac{L_p}{r_L}) \end{cases} \tag{5}$$

where $\mathrm{ceil}(\cdot)$ is round up to an integer and $\mathrm{rem}(\cdot)$ is the remainder. Then we would get a mapping list of $\mathbf{Z}_{[p,p+l_p^j]} \rightarrow \mathbf{U}_p$. The $r_L$ is set to 5.

Thirdly, the distance kernel matrix, $\mathbf{K} = \phi(\mathbf{U})^T \phi(\mathbf{U}) \in \mathrm{IR}^{m\times m}$, where $\mathbf{K} = \{\mathbf{k}_1, \mathbf{k}_2, ..., \mathbf{k}_M\}^T$ and $\mathbf{k}_i = (\kappa_i^1, \kappa_i^2, ..., \kappa_i^M)^T$ is calculated from $\mathbf{U}$. $\kappa_i^j$ defines the similarity between $\mathbf{U}_i$ and $\mathbf{U}_j$, where we use the Gaussian kernel [15] $\kappa_i^j = \exp\left[-\frac{(\mathbf{U}_i - \mathbf{U}_j)(\mathbf{U}_i - \mathbf{U}_j)^T}{2\sigma^2}\right]$. We set a fixed $\sigma$ as 30 to keep each $\mathbf{U}$ from different videos would have the same Gaussian kernel and map to the same high-dimensional space.

Fourthly, NMs are decomposed by aligned cluster analysis (ACA) [16,17] according to $\mathbf{K}$. ACA has two steps, initialization, and optimization. We use spectral clustering (SC) [18] for initialization. Let $\begin{bmatrix} \mathbf{D}_{11} & \cdots & \mathbf{D}_{1P} \\ \vdots & \ddots & \vdots \\ \mathbf{D}_{P1} & \cdots & \mathbf{D}_{PP} \end{bmatrix} = \mathbf{K}$ be the SC initialization distance kernel matrix, where each $\mathbf{D} = \{\mathbf{d}_1, \mathbf{d}_2, ..., \mathbf{d}_p\}^T$ is the initial NM matrix and $\mathbf{d} = (d_i^1, d_i^2, ..., d_i^p)^T$. We set the classes number $k_m$ of SC to 30. Then dynamic time alignment kernel (DTAK) is used as a metric for the optimization of ACA. Let $\mathbf{C}$ be the cumulative kernel matrix of $\mathbf{D}$, where $\mathbf{C}$ is defined as $\mathbf{C} = \{\mathbf{c}_1, \mathbf{c}_2, ..., \mathbf{c}_p\}^T$ and $\mathbf{c}$ is defined as $\mathbf{c} = (c_i^1, c_i^2, ..., c_i^p)^T$. DTAK uses $\mathbf{C}$ and dynamic programming to calculate the similarity of two time series as

$$\tau(\mathbf{c}_i, \mathbf{c}_j) = \frac{\mathbf{C}_p}{p_x + p_y}, c_i^j = \max \begin{cases} c_{i-1}^j + d_i^j \\ c_{i-1}^{j-1} + 2d_i^j \\ c_i^{j-1} + d_i^j \end{cases} \tag{6}$$

Then the energy function of ACA could be written as

$$J_{aca}(\mathbf{G},\mathbf{s})=\sum_{i=1}^{k_m}\sum_{j=1}^{P} g_{ij}\left\|\psi(\mathbf{V}^i_{[s_j,s_{j+1})})-\psi(\mathbf{z}_i)\right\|^2,\ \ n_{\min}\le s_{j+1}-s_j<n_{\max} \tag{7}$$

$$\left\|\psi(\mathbf{V}^i_{[s_j,s_{j+1})})-\psi(\mathbf{z}_i)\right\|^2=\tau_{jj}-\frac{2}{p_i}\sum_{q=1}^{p} g_{iq}\tau_{jq}+\frac{1}{p_i^2}\sum_{q_1,q_2=1}^{p} g_{iq_1}g_{iq_2}\tau_{q_1q_2} \tag{8}$$

where $\mathbf{G}\in\{0,1\}^{k_m\times P}$ is a class indicator matrix and $\mathbf{s}$ is a vector containing the start and end of each NM. The $g_{ij}$ is the element of $\mathbf{G}$ if $\mathbf{V}$ belongs to class $i$, $g_{ij}=1$, otherwise $g_{ij}=0$. The $\psi(\cdot)$ is a mapping of the sequence into a feature space. The $\mathbf{V}$ is a NM series to be optimized until optimal time duration. $\mathbf{z}$ is the center NM series of class $i$. $\left\|\psi(\mathbf{V}^i_{[s_j,s_{j+1})})-\psi(\mathbf{z}_i)\right\|^2$ represents the distance between two NM time series $\mathbf{V}$ and $\mathbf{z}$, which is calculated as (6), where $\tau$ is the DTAK of $\mathbf{V}$ and $\mathbf{z}$. In (5), the $[s_j,s_{j+1}]$ need a specific range, which means that the temporal scale of each NM is restricted. We set $n_{\min}$ to 100 ms and $n_{\max}$ to 2000 ms for containing a large temporal scale of NMs as far as possible. After determining all the parameters, the $J_{aca}$ could be optimized as

$$\mathbf{G},\mathbf{s}=\arg\min_{\mathbf{G},\mathbf{s}} J_{aca}(\mathbf{G},\mathbf{s})=\arg\min_{\mathbf{G},\mathbf{s}}\sum_{i=1}^{k_m}\sum_{j=1}^{P} g_{ij}\left\|\psi(\mathbf{V}^i_{[s_j,s_{j+1})})-\psi(\mathbf{z}_i)\right\|^2 \tag{9}$$

which could be converted to a recursive form:

$$J(\theta)=\min_{1<i\le\theta}\left(J(i-1)+\min_{\mathbf{G},\mathbf{s}} J_{aca}(\mathbf{G},\mathbf{s})\Big|_{\mathbf{V}_{[i,\theta]}}\right) \tag{10}$$

To optimize $\mathbf{G}$ and $\mathbf{s}$ faster, the DPSearch is used to reduce the computational cost of ACA, which is introduced in [16,17] in detail.

**The group segment kernel matrix and low dimensional embedding**

After the optimization of ACA, the optimal $\mathbf{G}$ and $\mathbf{s}$ are confirmed. Let $\mathbf{X}=\{\mathbf{x}_1,\mathbf{x}_2,...,\mathbf{x}_{39}\}^T$ be the NM time series and $\mathbf{x}_i=(x_i^1,x_i^2,...,x_i^N)^T$ contains the data points of NMs after alignment, spatial reduction, and temporal reduction. The optimal DTAK (Supplementary Fig. 7) distance matrix $\mathbf{T}$ could be calculated as

$$\mathbf{T}=\begin{pmatrix} tdist(\mathbf{X}_{s_1},\mathbf{X}_{s_1}) & \cdots & tdist(\mathbf{X}_{s_1},\mathbf{X}_{s_P}) \\ \vdots & \ddots & \vdots \\ tdist(\mathbf{X}_{s_P},\mathbf{X}_{s_1}) & \cdots & tdist(\mathbf{X}_{s_P},\mathbf{X}_{s_P}) \end{pmatrix} \tag{11}$$

where $tdist(\mathbf{X}_{s_i}, \mathbf{X}_{s_j}) = \left\| \psi(\mathbf{X}_{s_i}) - \psi(\mathbf{X}_{s_j}) \right\|^2$ is the DTAK distance between two NMs dynamic $\mathbf{X}_{s_i}$ and $\mathbf{X}_{s_j}$.

Then, the $\mathbf{T}$ is reduced dimensions from P dimensions to 2 dimensions by UMAP, which could make large amounts of data more widely distributed. Let $\mathbf{Y} = \{\mathbf{y}_1, \mathbf{y}_2\}^T$ be the two dimensions UMAP embedding of NMs, where each dimension $\mathbf{y}_i = (y_i^1, y_i^2, ..., y_i^P)^T$ contains the position of NMs in embedding. Then the velocity of each NM segment is added to embedding as the third dimension. The velocity dimension represents the locomotion of animals. Let $\mathbf{v} = (v_1, v_2, ..., v_P)^T$ be the velocity dimension and $\mathbf{C} = \{\mathbf{c}_1, \mathbf{c}_2, \mathbf{c}_3\}^T$ be the centroid of mice, then the mean velocity of each segment could be calculated as

$$v_i = \frac{Fs}{m_i} \left\| \Delta \mathbf{C}_{s_i} \right\|^2 \tag{12}$$

where $Fs{=}30$ is the framerate of each video, $m$ is the duration of the centroid segment $\mathbf{C}_s$ and $\Delta\mathbf{C}_s$ is the difference of $\mathbf{C}_s$. Velocity dimension is added as an orthogonal dimension to $\mathbf{Y}$ thus we get a 3D embedding $\mathbf{E} = \{\mathbf{y}_1, \mathbf{y}_2, \mathbf{v}\}^T$. The 3D embedding represents the patterns of NM and locomotion, which is built as a whole movement space of mice. To compare the three dimensions in the same unit, we get rid of the unit of them by Z-score in each dimension of $\mathbf{E}$. The $\mathbf{E}_z = \{\mathbf{z}_1, \mathbf{z}_2, \mathbf{z}_3\}^T$ denotes normalized embedding of $\mathbf{E}$ after Z-score [19].

**Unsupervised clustering**

Hierarchical clustering [20,21] is used to classify 3D embedding. The pairwise distance matrix $\mathbf{D}$ between $\mathbf{E}_z$ is calculated by a standardized Euclidean distance [22]. Then we use inner squared distance [23] to get the linkage of $\mathbf{D}$. The number of classes is set to 11 and 41 in a single video and multi-videos embedding clustering, respectively, which is used to cut the hierarchical cluster tree. Hence, we get the label of 3D embedding, the movements space, for further analysis of mice movements.

**Determining clustering number**

In the clustering task, selecting an appropriate number of clusters is important (Supplementary Fig. 10). However, in most cases, an appropriate number of clusters is difficult to choose because most of the movements have high similarity with each other, which causes the boundary of different movements clusters are blurry in movement space. Hence, we take careful consideration to determine the appropriate number of clusters by Bayesian Information Criterion (BIC) in 'mclust' package of R language [24]. That package provides 14 kinds of different models to infer the best parameters, such as the number of clusters. In our tests, the best numbers of clusters in single behavioral experiments are in the range of 10 to 20. Considering the discrimination of different behavior while giving them suitable labels, we choose 11 as the most appropriate number of clusters in movement clustering. When clustering all the movements of 10 *Shank3B*$^{+/+}$ mice and 10 *Shank3B*$^{-/-}$ mice in movement space, we use the recommended number of clusters of BIC with manual inspection, which is 41. The reason why we use 41 is that the number of clusters is too much to label all of them. Most of the movements have no differences in two mice groups, which means that we can only label the different movements and some critical movements.

**Behavioral phenotypes definition**

We define fourteen different kinds of behavioral phenotypes (Supplementary Table 1) referring to Mouse Ethogram data base (www.mousebehavior.org) and [25].

**Clustering quality index (CQI)**

To quantize the quality of clustering, we define CQI (Supplementary Fig. 10). Let DTAK distance matrix be $\mathbf{T} = [\mathbf{t}_1, \mathbf{t}_2, ..., \mathbf{t}_N]^T$ , and the movements features of the same cluster are $\mathbf{T}_m^{N_m} = [\mathbf{t}_m^1, \mathbf{t}_m^2, ..., \mathbf{t}_m^{N_m}]^T$ where $N_m$ is the number of $m$ movements. We first calculated the cross-correlation coefficient of intra and inter of clusters:

$$\begin{cases} \mathbf{c}_{\text{intra}} = xcorr(\mathbf{T}_m^{N_m^1}, \mathbf{T}_m^{N_m^2}) \\ \mathbf{c}_{\text{inter}} = xcorr(\mathbf{T}_m^{N_m^1}, \mathbf{T}^{N_m^3}) \end{cases} \tag{13}$$

where $\mathbf{c}_{\text{intra}} \in [-1,1]$ is the intra cross-correlation coefficient column vector of $m$ movements, $\mathbf{c}_{\text{inter}} \in [-1,1]$ is the inter cross-correlation coefficient column vector of $m$ movements, $\mathbf{T}_m^{N_m}$ is randomly divided into $\mathbf{T}_m^{N_m^1}$ and $\mathbf{T}_m^{N_m^2}$ in equal and $N_m^1 + N_m^2 = N_m$ , $\mathbf{T}_m^{N_m^3}$ is randomly selected from a complementary set of $\mathbf{T}_m^{N_m}$ and $\mathbf{T}_m^{N_m^1}$ has the same number of $\mathbf{T}_m^{N_m^3}$ . $\mathbf{c}_{\text{intra}}$ represents the similarity in a cluster of movements and $\mathbf{c}_{\text{inter}}$ represents the similarity between a cluster of movements with other clusters of movements. Then we use $\mathbf{c}_{\text{intra}}$ and $\mathbf{c}_{\text{inter}}$ to build a cartesian coordinate system $\mathbf{\Theta} = \{\mathbf{c}_{\text{inter}}, \mathbf{c}_{\text{intra}}\}$ . When the quality of clustering is much better, the value of $\mathbf{c}_{\text{intra}}$ should be close to $1$ , and the value of $\mathbf{c}_{\text{inter}}$ should be close to $-1$ . Thus, we could map $\mathbf{\Theta}$ to a new one-dimensional linear space:

$$\mathbf{\Phi} = \mathbf{A} \cdot [\mathbf{c}_{\text{intra}}, \mathbf{c}_{\text{inter}}]^T \tag{14}$$

where $\mathbf{A}=[-\frac{\sqrt{2}}{2},\frac{\sqrt{2}}{2}]$ is the projection matrix from $\mathbf{\Theta}$ to $\mathbf{\Phi}$. In $\mathbf{\Theta}$, $\mathbf{\Phi}$ could be regarded as a linear function $c_{\text{intra}}+c_{\text{inter}}=0$, which represents the relationships between $\mathbf{c}_{\text{intra}}$ and $\mathbf{c}_{\text{inter}}$ effectively. At last, we normalize $\mathbf{\Phi}$ to $[0,1]$ by sigmoid function to reduce the dynamic range of $\mathbf{\Phi}$ and $\text{CQI}=\text{sigmoid}(\mathbf{\Phi})$.

**Correlation analysis of clustering quality**

To compare the clustering similarity of inter and intra movement phenotype classes simultaneously, we applied the correlation analysis with linear regression for them [26]. The feature vector of each behavioral module ($1\times n$ vector) in DTAK distance matrix ($n\times n$) represents the distance between one behavioral module with each other, so that all the dimensions of feature vector have the same unit to be compared. If two behavioral modules have higher similarity, the correlation of their feature vector should be closed to each other. So, we choose one behavioral module's feature vector as the reference, and randomly select the feature vector of other behavioral modules as the targets for comparison. Each target with the reference forms a $2\times n$ list, which could be plotted on the two-dimensional space for linear regression and correlation analysis. The higher $R$ value of linear regression, the higher correlation of two behavioral modules.

For the inter class, we sequentially choose the behavioral module as reference, and randomly select three other behavioral modules' feature vector in the same class as targets to plot them on two dimensional space until all the references are plotted with targets. Then, we calculate the linear regression of them. For the intra class, we sequentially choose the behavioral module as reference, and randomly select three other behavioral modules' feature vector out of the reference's class then plot them on two dimensional space in gray points until all the references are plotted with targets. Similarly, we calculated the linear regression of them. It should be noted that only the regression line of inter classes are plotted on the figure.

**The mapping of new behavioral data to UMAP template**

If we directly construct low dimensional embedding by all the big behavioral data of 84 trails, it would consume a large number of unnecessary computing resource. To process the big behavioral data of all the groups, we map them to the UMAP template constructed by the behavioral modules of 6 *Shank3B*$^{+/+}$ *and 6 Shank3B*$^{-/-}$ mice. We use the UMAP toolbox of MATLAB and apply the default parameters to map the 84 trails of behavioral data to the template [1427]. After the mapping, the categories of new behavioral modules should be confirmed. We use K-Nearest-Neighbor (KNN) to assign categories for new behavioral modules in low dimensional embedding, and we use the default K for it [22].

**Moving intensity (MI)**

To quantize the movement of each body part in the body, we define a parameter of MI. We regard each body part as particle, which means that the moving intensity of it is correlated to velocity, and it accumulation with time. Further, the intensity of each movement segment should be the summation across time like the concept of energy. Therefore, we define MI like kinetic energy in physics, which is the energy possessed by an object under its motion,

$$E_k = \frac{1}{2} m v^2 \tag{15}$$

where $E_k$ is kinetic energy, $m$ is the mass of an object and $v$ is the velocity of an object. Considering that the body parts have unit mass and simplifying the calculation, we define $m_p = \frac{1}{2} m$ as the mass of body parts, and the value of $m_p$ is 1. Then the kinetic energy is transformed to MI and the MI of a movement in different coordinates could be calculated as

$$\begin{cases} E_{xy} = \dfrac{1}{2NT} \sum_{i=1}^{N} (\mathbf{V}_{x_i}^2 + \mathbf{V}_{y_i}^2) \\ E_{yz} = \dfrac{1}{2NT} \sum_{i=1}^{N} (\mathbf{V}_{y_i}^2 + \mathbf{V}_{z_i}^2) \\ E_{all} = \dfrac{1}{3NT} \sum_{i=1}^{N} (\mathbf{V}_{x_i}^2 + \mathbf{V}_{y_i}^2 + \mathbf{V}_{z_i}^2) \end{cases} \tag{16}$$

where $E_{xy}$ is the mean MI in XY coordinates plane, $E_{yz}$ is the mean MI in YZ coordinates plane, $E_{all}$ is the mean MI in XYZ coordinates space, $N$ is the dimension of movements, $\mathbf{V}$ is the velocity matrix of different movements and $T$ is the temporal length of $\mathbf{V}$.

After each MI of a body part in XY (horizontal) and YZ (vertical) plane coordinates were calculated, they were visualized in Fig. 5. We visualized them in specific top (XY) and side views (YZ). First, we averaged the positions of each body part across time to plot the average pose skeleton of each movement category, shown by solid lines. Second, we meshed the plane of the average pose skeleton to create grids for visualizing the MI. Each position of the skeleton has a corresponding MI parameter, which could be plotted on the grids. The MIs in the same grids were averaged to obtain the mean MI. Third, we calculated the frequency of representation of each body part in a grid cell to calculate a weighted MI. This step aims to make the position-correlated movement intensity of body parts describable by MI. Finally, the weighted MIs are plotted as heat map, in which it is easy to observe the movement areas of each body part and depict the movement intensity in specific positions.

**Traditional behavioral analysis**

To quantize the behaviors in a circular open field test, we first plot the velocity-trajectory maps for behavioral visualization then calculate mean velocity, maximum velocity, locomotion velocity [28] and mean anxiety index [29] of *Shank3B*$^{-/-}$ and *Shank3B*$^{+/+}$ mice (Supplementary Fig. 11). We extract the body part of back in the plane $\{\mathbf{X},\mathbf{Y}\}$ as the centroid of the mouse and apply a backward difference of centroids to calculate the instantaneous velocities of each frame. Then the instantaneous velocities of each frame are normalized to $[0,1]$ and they are plot at the positions of mouse centroids of each frame in the same canvas, which is the velocity-trajectory map after drawing all the points.

Mean velocity represents the mean activity of mice. Let mean velocity be $V_m$ , which could be calculated as

$$V_m = \frac{Fs}{N}\sum_{i=2}^{N}\sqrt{\left(x_c^i - x_c^{i-1}\right)^2 + \left(y_c^i - y_c^{i-1}\right)^2} \tag{17}$$

where $Fs{=}30$ is the framerate of each video, $N$ is the number of frames of each video and $(x_c^i, x_y^i)$ is the mouse centroid of each frame in the plane $\{\mathbf{X},\mathbf{Y}\}$. Let maximum velocity be $V_{\max}$ and instantaneous velocity be $V_i = Fs\sqrt{\left(x_c^i - x_c^{i-1}\right)^2 + \left(y_c^i - y_c^{i-1}\right)^2}$ , then $V_{\max}$ could be calculated as

$$V_{\max} = \max(\mathbf{V}) \tag{18}$$

where $\mathbf{V} = \left[V_2, V_3, \ldots, V_i\right]^T$ is the velocity vector. Let locomotion velocity be $V_l$, which could be calculated as

$$V_l = \frac{1}{M}\sum_{i=2}^{M} V_i, V_i > V_{\mathbf{thres}} \tag{19}$$

where $V_{\mathbf{thres}}$ is the threshold of $V_i$ to select the locomotion velocity from $V_i$ set as 190 and $M$ is the number of selected $V_i$. Mean anxiety index represents the anxious degree of mice. Let mean anxiety index be $A_m$, which could be calculated as

$$A_m = \frac{\sum\left[\mathbf{T}_m \wedge \left(\delta\left(x_{tc}, y_{tc}\right) * \mathbf{S}_{\frac{R}{2}}\right)\right]}{\sum \mathbf{T}_m} \tag{20}$$

where $\mathbf{T}_m$ is the trajectory maps of each mouse, $\delta$ is the 2D impulse function with the same size of $\mathbf{T}_m$, $(x_{tc}, y_{tc})$ is the center of $\mathbf{T}_m$ which makes the impulse of $\delta$ locate to the center of $\mathbf{T}_m$, $R$ is the radius of circular open field, $\mathbf{S}_{\frac{R}{2}}$ is a disk structural element with the half radius of circular open field, $\delta\left(x_{tc}, y_{tc}\right) * \mathbf{S}_{\frac{R}{2}}$ is the 2D convolution of $\delta$ and $\mathbf{S}_{\frac{R}{2}}$ which sets the disk structural element at the center of $\mathbf{T}_m$, $\mathbf{T}_m \wedge \left(\delta\left(x_{tc}, y_{tc}\right) * \mathbf{S}_{\frac{R}{2}}\right)$ means extracting $\mathbf{T}_m$ of the central circular region in the range of $\mathbf{S}_{\frac{R}{2}}$ by AND operation and $\sum(\cdot)$ sums over the values of the matrix elements.

**Statistics**

Analyses are performed using Prism 8.0 (GraphPad Software). Before hypothesis testing, data are first tested for normality by the Shapiro-Wilk normality test and tested for homoscedasticity by F test. If the null hypothesis that the data comes from a normal distribution cannot be rejected and the null hypothesis that the data have homogeneity of variances cannot be rejected, parametric tests are used (for example, Student's t-test for two groups, two-way ANOVA with Holm–Sidak post-hoc test for more than two groups). If the null hypothesis that the data comes from a normal distribution can be rejected and the null hypothesis that the data have homogeneity of variances can be rejected, nonparametric tests are used (for example, Mann Whitney test for two groups, two-way ANOVA with Bonferroni's test for more than two groups).

The data of mean velocity of 6 *Shank3B*$^{+/+}$ mice group and 6 *Shank3B*$^{-/-}$ mice group (fig.5 B) are normality and have homogeneity of variances so that we use unpaired t-test to compare the difference between them. The data of the mean anxiety index of two groups (fig.5 B) are non-normality, so that we use the Mann-Whitney test to compare the difference between them. The data of movements' fractions (fig.5 D) are normality and have homogeneity of variances so that we use two-way ANOVA with Holm–Sidak post-hoc test to compare the difference between each group of them.

**Autistic-like behavior space**

The proportion of 41 clusters movements of each mouse (fig. 6 D) could be regarded as a feature vector of mouse behavior space. Let the feature vector be $\mathbf{x} = [x_1, x_2, ..., x_{41}]^T$, the feature matrix of behavior space is $\mathbf{X} = [\mathbf{x}_1, \mathbf{x}_2, ..., \mathbf{x}_{12}]^T$. Then we use UMAP to reduce the feature dimensions of $\mathbf{X}$ from 41 to 3,

$$\mathbf{Y} = f_{UMAP}(\mathbf{X}) \tag{21}$$

where $\mathbf{Y} = [\mathbf{y}_1, \mathbf{y}_2, \mathbf{y}_3]^T$, $\mathbf{y}_i = [y_i^1, y_i^2, ..., y_i^{41}]^T$ is the 3D feature matrix, the autistic-like behavior space, after the dimension reduction of UMAP. $f_{UMAP}(\cdot)$ have the parameters n_neighbors to 30 and min_dist to 0.3, which are robust enough to change in a wide range and keep the discrimination of *Shank3B*$^{+/+}$ mice group and *Shank3B*$^{-/-}$ mice group in autistic-like behavior space. To quantize the discrimination of two groups, we fit a linear classification model to $\mathbf{Y}$ by the fitclinear function in MATLAB with default parameters.

**The analysis of state transitions**

In the behavioral analysis of continuous long-term monitoring, we applied state transitions analysis for behavioral modules (Supplementary Fig. 13 e, f, g) [303132]. The 31 behavioral modules are regarded as states in probabilistic graphical model especially. The probabilities of states are calculated approximately by the proportion of the number of behavioral modules. The states transition probabilities are calculated approximately by the transition proportion of the number of previous behavioral modules to current behavioral modules. The difference value of transition probabilities between Night and Daytime are calculated approximately by the subtraction the behavioral modules' proportion of Daytime from Night.

#### Supplementary Tables

**Supplementary Table 1 Definition of the behaviors (ethogram) for manual labeling**

| Behavior | Definition |
|---|---|
| Running | The mouse locomotes with relatively high speed. |
| Trotting | The mouse locomotes at a slow and intermittent pace. |
| Stepping | The mouse takes a step forward with a short distance locomotion. |
| Diving | The mouse bends down its body from standing state then stretches forward. |
| Sniffing | The mouse investigates environment with the nose held in the air or contacts the environment with nose closely. |
| Rising | The mouse rises from four legs on the ground to steadily stand on its hind legs. |
| Right turning | The mouse bends its body to right or turns body to right while walking. |
| Up stretching | The mouse stands on its hind legs while the front part of body stretches back and forth. |
| Falling | The mouse takes its croup as pivot and gets down with straight back from standing on its hind legs. |
| Left turning | The mouse bends its body to left or turns body to left while walking. |
| Walking | The mouse locomotes with relatively low speed. |
| Rearing | The mouse stands on its hind legs, and the back is straight. |
| Hunching | The mouse stands on its hind legs while the back is bent. |
| Self-grooming | The mouse licks its fur, grooms with the forepaws, or scratches with any limb. |

### Supplementary Table 2 The SHANK3 mice used in group comparison

| Experiments index | Date | Start time | Mice index | Genotypes | Ages (weeks) | Sexes | Environments | Light conditions |
|---|---|---|---|---|---|---|---|---|
| 1 | 20201224 | 13:41 | 1030 | Shank3B$^{-/-}$ | 5-6 | Male | Circular open field | White light |
| 2 | 20201224 | 15:33 | 1038 | Shank3B$^{+/+}$ | 5-6 | Male | Circular open field | White light |
| 3 | 20201224 | 16:27 | 1042 | Shank3B$^{+/+}$ | 5-6 | Male | Circular open field | White light |
| 4 | 20201224 | 17:20 | 1051 | Shank3B$^{+/+}$ | 5-6 | Male | Circular open field | White light |
| 5 | 20201225 | 12:05 | 765 | Shank3B$^{+/+}$ | 11-13 | Male | Circular open field | White light |
| 6 | 20201225 | 12:33 | 771 | Shank3B$^{-/-}$ | 11-13 | Male | Circular open field | White light |
| 7 | 20201225 | 13:00 | 782 | Shank3B$^{-/-}$ | 11-13 | Male | Circular open field | White light |
| 8 | 20201225 | 13:27 | 789 | Shank3B$^{-/-}$ | 11-13 | Male | Circular open field | White light |
| 9 | 20201225 | 13:54 | 810 | Shank3B$^{+/+}$ | 11-13 | Male | Circular open field | White light |
| 10 | 20201225 | 14:22 | 812 | Shank3B$^{+/+}$ | 11-13 | Male | Circular open field | White light |
| 11 | 20201225 | 14:49 | 814 | Shank3B$^{-/-}$ | 11-13 | Male | Circular open field | White light |
| 12 | 20201225 | 15:16 | 818 | Shank3B$^{+/+}$ | 11-13 | Male | Circular open field | White light |
| 13 | 20201225 | 16:12 | 753 | Shank3B$^{-/-}$ | 11-13 | Male | Circular open field | White light |
| 14 | 20201225 | 17:08 | 1053 | Shank3B$^{+/+}$ | 5-6 | Male | Circular open field | White light |
| 15 | 20201226 | 12:25 | 754 | Shank3B$^{+/+}$ | 11-13 | Male | Circular open field | White light |
| 16 | 20201226 | 13:49 | 829 | Shank3B$^{-/-}$ | 11-13 | Male | Circular open field | White light |
| 17 | 20201226 | 14:17 | 985 | Shank3B$^{+/+}$ | 11-13 | Male | Circular open field | White light |
| 18 | 20201226 | 14:46 | 986 | Shank3B$^{-/-}$ | 11-13 | Male | Circular open field | White light |
| 19 | 20201226 | 15:13 | 982 | Shank3B$^{+/+}$ | 11-13 | Male | Circular open field | White light |
| 20 | 20201226 | 15:41 | 767 | Shank3B$^{+/+}$ | 11-13 | Male | Circular open field | White light |
| 21 | 20201226 | 16:36 | 773 | Shank3B$^{-/-}$ | 11-13 | Male | Circular open field | White light |
| 22 | 20201227 | 12:08 | 1032 | Shank3B$^{+/+}$ | 5-6 | Female | Circular open field | White light |
| 23 | 20201227 | 12:36 | 1036 | Shank3B$^{+/+}$ | 5-6 | Female | Circular open field | White light |
| 24 | 20201227 | 13:32 | 1050 | Shank3B$^{-/-}$ | 5-6 | Female | Circular open field | White light |
| 25 | 20201227 | 14:00 | 1068 | Shank3B$^{-/-}$ | 5-6 | Female | Circular open field | White light |
| 26 | 20201227 | 14:26 | 1061 | Shank3B$^{+/+}$ | 5-6 | Female | Circular open field | White light |
| 27 | 20201227 | 14:53 | 1062 | Shank3B$^{+/+}$ | 5-6 | Female | Circular open field | White light |
| 28 | 20201227 | 15:21 | 1063 | Shank3B$^{-/-}$ | 5-6 | Female | Circular open field | White light |
| 29 | 20201227 | 16:16 | 1065 | Shank3B$^{+/+}$ | 5-6 | Female | Circular open field | White light |
| 30 | 20201228 | 11:57 | 1029 | Shank3B$^{+/+}$ | 5-6 | Male | Square open field | White light |
| 31 | 20201228 | 12:26 | 1030 | Shank3B$^{-/-}$ | 5-6 | Male | Square open field | White light |
| 32 | 20201228 | 12:53 | 1031 | Shank3B$^{-/-}$ | 5-6 | Male | Square open field | White light |
| 33 | 20201228 | 13:21 | 1033 | Shank3B$^{+/+}$ | 5-6 | Male | Square open field | White light |
| 34 | 20201228 | 13:49 | 1035 | Shank3B$^{+/+}$ | 5-6 | Male | Square open field | White light |
| 35 | 20201228 | 14:18 | 1038 | Shank3B$^{+/+}$ | 5-6 | Male | Square open field | White light |
| 36 | 20201228 | 14:46 | 1040 | Shank3B$^{-/-}$ | 5-6 | Male | Square open field | White light |
| 37 | 20201228 | 15:14 | 1042 | Shank3B$^{+/+}$ | 5-6 | Male | Square open field | White light |
| 38 | 20201228 | 15:41 | 1046 | Shank3B$^{+/+}$ | 5-6 | Male | Square open field | White light |
| 39 | 20201228 | 16:08 | 1051 | Shank3B$^{+/+}$ | 5-6 | Male | Square open field | White light |
| 40 | 20201228 | 17:02 | 1053 | Shank3B$^{+/+}$ | 5-6 | Male | Square open field | White light |
| 41 | 20201229 | 14:04 | 1029 | Shank3B$^{+/+}$ | 5-6 | Male | Circular open field | Infrared |
| 42 | 20201229 | 14:31 | 1030 | Shank3B$^{-/-}$ | 5-6 | Male | Circular open field | Infrared |
| 43 | 20201229 | 14:59 | 1031 | Shank3B$^{-/-}$ | 5-6 | Male | Circular open field | Infrared |
| 44 | 20201229 | 15:26 | 1033 | Shank3B$^{+/+}$ | 5-6 | Male | Circular open field | Infrared |
| 45 | 20201229 | 15:54 | 1051 | Shank3B$^{+/+}$ | 5-6 | Male | Circular open field | Infrared |
| 46 | 20201229 | 16:48 | 1053 | Shank3B$^{+/+}$ | 5-6 | Male | Circular open field | Infrared |
| 47 | 20201229 | 17:15 | 1035 | Shank3B$^{+/+}$ | 5-6 | Male | Circular open field | Infrared |

| | | | | | | | | |
|---|---|---|---|---|---|---|---|---|
| 48 | 20201229 | 15:42 | 1038 | Shank3B$^{+/+}$ | 5-6 | Male | Circular open field | Infrared |
| 49 | 20201229 | 18:10 | 1040 | Shank3B$^{-/-}$ | 5-6 | Male | Circular open field | Infrared |
| 50 | 20201229 | 18:38 | 1042 | Shank3B$^{+/+}$ | 5-6 | Male | Circular open field | Infrared |
| 51 | 20201229 | 19:03 | 1046 | Shank3B$^{+/+}$ | 5-6 | Male | Circular open field | Infrared |
| 52 | 20201230 | 11:47 | 69 | Shank3B$^{-/-}$ | 5-6 | Male | Circular open field | White light |
| 53 | 20201230 | 12:44 | 65 | Shank3B$^{-/-}$ | 5-6 | Male | Circular open field | White light |
| 54 | 20201230 | 13:12 | 12 | Shank3B$^{-/-}$ | 5-6 | Male | Circular open field | White light |
| 55 | 20201230 | 14:36 | 58 | Shank3B$^{-/-}$ | 5-6 | Male | Circular open field | White light |
| 56 | 20201230 | 15:07 | 69 | Shank3B$^{-/-}$ | 5-6 | Male | Square open field | White light |
| 57 | 20201231 | 11:43 | 65 | Shank3B$^{-/-}$ | 5-6 | Male | Circular open field | Infrared |
| 58 | 20201231 | 12:11 | 12 | Shank3B$^{-/-}$ | 5-6 | Male | Circular open field | Infrared |
| 59 | 20201231 | 13:04 | 57 | Shank3B$^{-/-}$ | 5-6 | Male | Circular open field | Infrared |
| 60 | 20201231 | 13:32 | 58 | Shank3B$^{-/-}$ | 5-6 | Male | Circular open field | Infrared |
| 61 | 20201231 | 14:47 | 12 | Shank3B$^{-/-}$ | 5-6 | Male | Square open field | White light |
| 62 | 20201231 | 15:41 | 57 | Shank3B$^{-/-}$ | 5-6 | Male | Square open field | White light |
| 63 | 20201231 | 16:09 | 58 | Shank3B$^{-/-}$ | 5-6 | Male | Square open field | White light |
| 64 | 20201231 | 16:40 | 69 | Shank3B$^{-/-}$ | 5-6 | Male | Circular open field | Infrared |
| 65 | 20210101 | 12:20 | 22 | Shank3B$^{-/-}$ | 5-6 | Female | Circular open field | White light |
| 66 | 20210101 | 12:48 | 24 | Shank3B$^{+/+}$ | 5-6 | Female | Circular open field | White light |
| 67 | 20210101 | 13:15 | 25 | Shank3B$^{+/+}$ | 5-6 | Female | Circular open field | White light |
| 68 | 20210101 | 13:41 | 19 | Shank3B$^{-/-}$ | 5-6 | Female | Circular open field | White light |
| 69 | 20210101 | 14:36 | 40 | Shank3B$^{-/-}$ | 5-6 | Female | Circular open field | White light |
| 70 | 20210101 | 15:08 | 33 | Shank3B$^{-/-}$ | 5-6 | Female | Circular open field | White light |
| 71 | 20210101 | 15:35 | 41 | Shank3B$^{-/-}$ | 5-6 | Female | Circular open field | White light |
| 72 | 20210101 | 14:02 | 45 | Shank3B$^{+/+}$ | 5-6 | Female | Circular open field | White light |
| 73 | 20210107 | 20:20 | H1 | C57BL/6J | 13 | Male | Circular home-cage | Infrared cameras (12 hours white light and 12 hours dark) |
| 74 | 20200623 | 10:40 | A61 | Shank3B$^{+/+}$ | 5-6 | Male | Circular open field | White light |
| 75 | 20200623 | 11:07 | A62 | Shank3B$^{+/+}$ | 5-6 | Male | Circular open field | White light |
| 76 | 20200623 | 11:35 | A63 | Shank3B$^{-/-}$ | 5-6 | Male | Circular open field | White light |
| 77 | 20200623 | 12:03 | A64 | Shank3B$^{-/-}$ | 5-6 | Male | Circular open field | White light |
| 78 | 20200623 | 12:32 | A71 | Shank3B$^{+/+}$ | 5-6 | Male | Circular open field | White light |
| 79 | 20200623 | 12:59 | A72 | Shank3B$^{+/+}$ | 5-6 | Male | Circular open field | White light |
| 80 | 20200623 | 14:33 | A73 | Shank3B$^{-/-}$ | 5-6 | Male | Circular open field | White light |
| 81 | 20200623 | 15:01 | A74 | Shank3B$^{-/-}$ | 5-6 | Male | Circular open field | White light |
| 82 | 20200623 | 15:31 | A75 | Shank3B$^{-/-}$ | 5-6 | Male | Circular open field | White light |
| 83 | 20200623 | 15:58 | A76 | Shank3B$^{-/-}$ | 5-6 | Male | Circular open field | White light |
| 84 | 20200623 | 16:26 | G81 | Shank3B$^{+/+}$ | 5-6 | Male | Circular open field | White light |
| 85 | 20200623 | 16:53 | G82 | Shank3B$^{+/+}$ | 5-6 | Male | Circular open field | White light |

**Supplementary Table 3 The individual level comparison between *Shank3B*$^{+/+}$ and *Shank3B*$^{-/-}$ mice**

| | KO1 | KO2 | KO3 | KO4 | KO5 | KO6 | KO7 | KO8 | KO9 | KO10 | Group Difference ↑ |
|---|---|---|---|---|---|---|---|---|---|---|---|
| **WT1** | M5↓ | M5↓ | M5↓ | M5↓ | M5↓ | M5↓ | M5↓ | M5↓ | M5↓ | M5↓ | M5: 0 |
| | M14↓ | M14↓ | M14↓ | M14↓ | M14↓ | M14 | M14↓ | M14↓ | M14↓ | M14↑ | M14: 2 |
| | M21↓ | M21↓ | M21↓ | M21↓ | M21↓ | M21↓ | M21↓ | M21↓ | M21↓ | M21↓ | M21: 0 |
| | M22↓ | M22↓ | M22↓ | M22↓ | M22↓ | M22↓ | M22↓ | M22↓ | M22↓ | M22↓ | M22: 0 |
| | M38↑ | M38↑ | M38↑ | M38↑ | M38↑ | M38↑ | M38↑ | M38↑ | M38↑ | M38↑ | M38: 10 |
| | M39↑ | M39↑ | M39↑ | M39↑ | M39↑ | M39↑ | M39↓ | M39↑ | M39↑ | M39↑ | M39: 9 |
| | M40↑ | M40↑ | M40↑ | M40↑ | M40↑ | M40↑ | M40↑ | M40↑ | M40↑ | M40↑ | M40: 10 |
| | M41↑ | M41↑ | M41↑ | M41↑ | M41↑ | M41↑ | M41↑ | M41↑ | M41↑ | M41↑ | M41: 10 |
| **WT2** | M5↓ | M5↓ | M5↓ | M5↓ | M5↓ | M5↓ | M5↓ | M5↓ | M5↓ | M5↓ | M5: 0 |
| | M14↓ | M14↓ | M14↓ | M14↓ | M14↓ | M14↓ | M14↓ | M14↓ | M14↓ | M14↓ | M14: 0 |
| | M21↓ | M21↓ | M21↓ | M21↓ | M21↓ | M21↓ | M21↓ | M21↓ | M21↓ | M21↓ | M21: 0 |
| | M22↓ | M22↓ | M22↓ | M22↓ | M22↓ | M22↓ | M22↓ | M22↑ | M22↓ | M22↓ | M22: 1 |
| | M38↑ | M38↑ | M38↑ | M38↑ | M38↑ | M38↓ | M38↓ | M38↑ | M38↑ | M38↑ | M38: 8 |
| | M39↑ | M39↑ | M39↑ | M39↑ | M39↑ | M39↑ | M39↑ | M39↑ | M39↑ | M39↑ | M39: 10 |
| | M40↑ | M40↑ | M40↑ | M40↑ | M40↑ | M40↑ | M40↑ | M40↑ | M40↑ | M40↑ | M40: 10 |
| | M41↑ | M41↑ | M41↑ | M41↑ | M41↑ | M41↑ | M41↑ | M41↑ | M41↑ | M41↑ | M41: 10 |
| **WT3** | M5↓ | M5↓ | M5↓ | M5↓ | M5↓ | M5↓ | M5↓ | M5↓ | M5↓ | M5↓ | M5: 0 |
| | M14↓ | M14↓ | M14↓ | M14↓ | M14↓ | M14↓ | M14↓ | M14↓ | M14↓ | M14↓ | M14: 0 |
| | M21↓ | M21↓ | M21↓ | M21↓ | M21↓ | M21↑ | M21↓ | M21↓ | M21↓ | M21↓ | M21: 1 |
| | M22↓ | M22↓ | M22↓ | M22↓ | M22↓ | M22↓ | M22↓ | M22↓ | M22↓ | M22↓ | M22: 0 |
| | M38↑ | M38↑ | M38↑ | M38↑ | M38↑ | M38↑ | M38↓ | M38↑ | M38↑ | M38↑ | M38: 8 |
| | M39↑ | M39↑ | M39↑ | M39↑ | M39↑ | M39↑ | M39↓ | M39↑ | M39↑ | M39↑ | M39: 9 |
| | M40↑ | M40↑ | M40↑ | M40↑ | M40↑ | M40↑ | M40↑ | M40↑ | M40↑ | M40↑ | M40: 10 |
| | M41↑ | M41↑ | M41↑ | M41↑ | M41↑ | M41↑ | M41↓ | M41↑ | M41↑ | M41↑ | M41: 9 |
| **WT4** | M5↓ | M5↓ | M5↓ | M5↓ | M5↓ | M5↓ | M5↓ | M5↓ | M5↓ | M5↓ | M5: 0 |
| | M14↓ | M14↓ | M14↓ | M14↓ | M14↓ | M14↓ | M14↓ | M14↓ | M14↓ | M14↑ | M14: 1 |
| | M21↓ | M21↓ | M21↓ | M21↓ | M21↓ | M21↓ | M21↓ | M21↓ | M21↓ | M21↓ | M21: 0 |
| | M22↓ | M22↓ | M22↓ | M22↓ | M22↓ | M22↑ | M22↑ | M22↑ | M22↓ | M22↓ | M22: 3 |
| | M38↑ | M38↑ | M38↑ | M38↑ | M38↑ | M38↑ | M38↑ | M38↑ | M38↑ | M38↑ | M38: 10 |
| | M39↑ | M39↑ | M39↑ | M39↑ | M39↑ | M39↑ | M39↑ | M39↑ | M39↑ | M39↑ | M39: 10 |
| | M40↑ | M40↑ | M40↑ | M40↑ | M40↑ | M40↑ | M40↑ | M40↑ | M40↑ | M40↑ | M40: 10 |
| | M41↑ | M41↑ | M41↑ | M41↑ | M41↑ | M41↑ | M41↑ | M41↑ | M41↑ | M41↑ | M41: 10 |
| **WT5** | M5↓ | M5↓ | M5↓ | M5↓ | M5↓ | M5↓ | M5↓ | M5↓ | M5↓ | M5↓ | M5: 0 |
| | M14↓ | M14↓ | M14↓ | M14↓ | M14↓ | M14↓ | M14↓ | M14↓ | M14↓ | M14↓ | M14: 0 |
| | M21↓ | M21↓ | M21↓ | M21↓ | M21↓ | M21↑ | M21↑ | M21↑ | M21↑ | M21↑ | M21: 5 |
| | M22↓ | M22↓ | M22↓ | M22↓ | M22↓ | M22↑ | M22↑ | M22↑ | M22↓ | M22↓ | M22: 3 |
| | M38↑ | M38↑ | M38↑ | M38↑ | M38↑ | M38↑ | M38↑ | M38↑ | M38↑ | M38↑ | M38: 10 |
| | M39↑ | M39↑ | M39↑ | M39↑ | M39↑ | M39↑ | M39↓ | M39↑ | M39↑ | M39↑ | M39: 9 |
| | M40↑ | M40↑ | M40↑ | M40↑ | M40↑ | M40↑ | M40↓ | M40↑ | M40↑ | M40↑ | M40: 9 |
| | M41↑ | M41↑ | M41↑ | M41↑ | M41↑ | M41↑ | M41↓ | M41↑ | M41↑ | M41↑ | M41: 9 |

| | | | | | | | | | | | |
|---|---|---|---|---|---|---|---|---|---|---|---|
| **WT6** | M5↓ | M5↓ | M5↓ | M5↓ | M5↑ | M5↓ | M5↓ | M5↓ | M5↑ | M5↓ | M5: 2 |
| | M14↓ | M14↓ | M14↓ | M14↓ | M14↑ | M14↑ | M14↓ | M14↑ | M14↓ | M14↑ | M14: 4 |
| | M21↓ | M21↓ | M21↓ | M21↓ | M21↓ | M21↑ | M21↓ | M21↑ | M21↑ | M21↓ | M21: 3 |
| | M22↓ | M22↓ | M22↓ | M22↓ | M22↓ | M22↓ | M22↓ | M22↑ | M22↓ | M22↓ | M22: 1 |
| | M38↑ | M38↑ | M38↑ | M38↑ | M38↑ | M38↑ | M38↑ | M38↑ | M38↑ | M38↑ | M38: 10 |
| | M39↑ | M39↑ | M39↑ | M39↑ | M39↑ | M39↑ | M39↑ | M39↑ | M39↑ | M39↑ | M39: 10 |
| | M40↑ | M40↑ | M40↑ | M40↑ | M40↑ | M40↑ | M40↓ | M40↑ | M40↓ | M40↑ | M40: 8 |
| | M41↑ | M41↑ | M41↑ | M41↑ | M41↑ | M41↑ | M41↓ | M41↓ | M41↑ | M41↑ | M41: 8 |
| **WT7** | M5↓ | M5↓ | M5↓ | M5↓ | M5↓ | M5↓ | M5↓ | M5↓ | M5↓ | M5↓ | M5: 0 |
| | M14↓ | M14↓ | M14↓ | M14↓ | M14↓ | M14↑ | M14↓ | M14↓ | M14↓ | M14↑ | M14: 2 |
| | M21↓ | M21↓ | M21↓ | M21↓ | M21↓ | M21↑ | M21↓ | M21↓ | M21↓ | M21↓ | M21: 1 |
| | M22↓ | M22↓ | M22↓ | M22↓ | M22↓ | M22↓ | M22↓ | M22↓ | M22↓ | M22↓ | M22: 0 |
| | M38↑ | M38↑ | M38↑ | M38↑ | M38↑ | M38↓ | M38↓ | M38↑ | M38↑ | M38↑ | M38: 8 |
| | M39↑ | M39↑ | M39↑ | M39↑ | M39↑ | M39↑ | M39↓ | M39↑ | M39↑ | M39↑ | M39: 9 |
| | M40↑ | M40↑ | M40↑ | M40↑ | M40↑ | M40↑ | M40↓ | M40↑ | M40↑ | M40↑ | M40: 9 |
| | M41↑ | M41↑ | M41↑ | M41↑ | M41↑ | M41↑ | M41↑ | M41↑ | M41↑ | M41↑ | M41: 10 |
| **WT8** | M5↓ | M5↑ | M5↓ | M5↓ | M5↑ | M5↑ | M5↑ | M5↓ | M5↑ | M5↑ | M5: 6 |
| | M14↓ | M14↓ | M14↓ | M14↓ | M14↑ | M14↑ | M14↓ | M14↑ | M14↓ | M14↑ | M14: 4 |
| | M21↓ | M21↓ | M21↓ | M21↓ | M21↓ | M21↑ | M21↓ | M21↓ | M21↓ | M21↓ | M21: 1 |
| | M22↓ | M22↓ | M22↓ | M22↓ | M22↓ | M22↑ | M22↓ | M22↑ | M22↓ | M22↓ | M22: 3 |
| | M38↑ | M38↑ | M38↑ | M38↑ | M38↑ | M38↓ | M38↓ | M38↑ | M38↓ | M38↑ | M38: 7 |
| | M39↑ | M39↑ | M39↑ | M39↑ | M39↑ | M39↓ | M39↓ | M39↑ | M39↑ | M39↑ | M39: 8 |
| | M40↑ | M40↑ | M40↑ | M40↑ | M40↑ | M40↑ | M40↓ | M40↑ | M40↑ | M40↑ | M40: 9 |
| | M41↑ | M41↑ | M41↑ | M41↑ | M41↑ | M41↑ | M41↑ | M41↑ | M41↑ | M41↑ | M41: 10 |
| **WT9** | M5↓ | M5↓ | M5↓ | M5↓ | M5↓ | M5↓ | M5↓ | M5↓ | M5↓ | M5↓ | M5: 0 |
| | M14↓ | M14↓ | M14↓ | M14↓ | M14↓ | M14↓ | M14↓ | M14↓ | M14↓ | M14↑ | M14: 1 |
| | M21↓ | M21↓ | M21↓ | M21↓ | M21↓ | M21↑ | M21↓ | M21↓ | M21↓ | M21↓ | M21: 1 |
| | M22↓ | M22↓ | M22↓ | M22↓ | M22↓ | M22↑ | M22↑ | M22↑ | M22↓ | M22↓ | M22: 3 |
| | M38↑ | M38↑ | M38↑ | M38↑ | M38↑ | M38↓ | M38↓ | M38↑ | M38↓ | M38↑ | M38: 7 |
| | M39↑ | M39↑ | M39↑ | M39↑ | M39↑ | M39↑ | M39↓ | M39↑ | M39↑ | M39↑ | M39: 9 |
| | M40↑ | M40↑ | M40↑ | M40↑ | M40↑ | M40↑ | M40↑ | M40↑ | M40↑ | M40↑ | M40: 10 |
| | M41↑ | M41↑ | M41↑ | M41↑ | M41↑ | M41↑ | M41↑ | M41↑ | M41↑ | M41↑ | M41: 10 |
| **WT10** | M5↓ | M5↓ | M5↓ | M5↓ | M5↓ | M5↓ | M5↓ | M5↓ | M5↓ | M5↓ | M5: 0 |
| | M14↓ | M14↓ | M14↓ | M14↓ | M14↓ | M14↓ | M14↓ | M14↓ | M14↓ | M14↓ | M14: 0 |
| | M21↓ | M21↓ | M21↓ | M21↓ | M21↓ | M21↑ | M21↓ | M21↓ | M21↓ | M21↓ | M21: 1 |
| | M22↓ | M22↓ | M22↓ | M22↓ | M22↓ | M22↓ | M22↓ | M22↓ | M22↓ | M22↓ | M22: 0 |
| | M38↑ | M38↑ | M38↑ | M38↑ | M38↑ | M38↑ | M38↓ | M38↑ | M38↑ | M38↑ | M38: 9 |
| | M39↑ | M39↑ | M39↑ | M39↑ | M39↑ | M39↑ | M39↑ | M39↑ | M39↑ | M39↑ | M39: 10 |
| | M40↑ | M40↑ | M40↑ | M40↑ | M40↑ | M40↑ | M40↓ | M40↑ | M40↑ | M40↑ | M40: 9 |
| | M41↑ | M41↑ | M41↑ | M41↑ | M41↑ | M41↑ | M41↑ | M41↑ | M41↑ | M41↑ | M41: 10 |
| **All Group Difference↑** | M5: 8, M14: 14, M21: 13, M22: 14, M38: 87, M39: 93, M40: 94, M41: 96 | | | | | | | | | | |

M*X*: *X* the index of behavioral module;

↑: KO individual shows higher fraction of *X* behavioral module than WT;

↓: KO individual shows lower fraction of *X* behavioral module than WT;

Group Difference ↑(M*X*: *n*): The number of KO individuals with higher *X* behavior modulus fractions than WT is n;

All Group Difference ↑(MX: *m*): In all the 100 times paired-wise comparisons, the number of times that KO individuals with higher *X* behavior modulus fractions than WT is m.
